# Ultrastructure of Inputs to the Granule Cell Domain of the Dorsal Cochlear Nucleus

**DOI:** 10.64898/2025.12.14.694253

**Authors:** Xiping Zhan, Aneesha Penn, Christina Rosette, Gadir Kaki, David K. Ryugo

## Abstract

The granule cell domain of the cochlear nucleus (GCD) receives input from multimodal brain regions including the second cervical spinal ganglion, the spinal trigeminal nucleus, the cuneate nucleus, and the lateral reticular nucleus. Most of the input are in the form of large mossy fibers and small boutons containing round synaptic vesicles and forming asymmetric synapses. A smaller number of inputs arise from the lateral reticular nucleus and the spinal trigeminal nucleus that contain pleomorphic synaptic vesicles implying inhibitory action. This circumstance positions the GCD for integrating polysensory excitatory and inhibitory inputs at the earliest stages of auditory processing, which suggests a role for segregating sound streams. The structural substrate for such a function is naturally elaborate. Mossy fibers from these different origins are large, multilobed endings that form synaptic glomeruli with postsynaptic targets that include granule cells, unipolar brush cells, cartwheel cells, and chestnut cells, which in turn project to principal cells in the dorsal cochlear nucleus (DCN). Numerous synapses contribute to the integration of the different modalities, but the organization of these synapses is still not well understood. We investigated this issue by labelling the presynaptic endings of different somatosensory inputs using biotinylated dextran amine or Phaseolus vulgaris leucoagglutinin, and studying their post-synaptic relationships in the GCD. The anterogradely-labeled endings and associated targets were visualized by light and electron microscopy. Following computer-aided, three-dimensional reconstructions, we found that unipolar brush cells are the main target of the C2 DRG and cuneate nucleus. Mossy fibers from the spinal trigeminal nucleus project to Golgi cells, an inhibitory neuron. In contrast, inhibitory boutons originated from the lateral reticular nucleus. These results reveal that somatosensory integration in the granule cell domain is achieved by distinctive synaptic wiring and specialized chemical signaling.

## Introduction

Sound is encoded by the auditory system but gains a broader context by the inclusion of additional systems. In mammals, a variety of evidence supports the idea of polysensory integration at multiple points along the auditory system that incorporates sensorimotor, learning, memory and emotional influences to facilitate the sorting of sound sources. The dorsal cochlear nucleus (DCN) is the first nucleus in the central auditory pathway to show such integration (Burian and Gstoettner, 1988; Kevetter and Perachio, 1989; Ryugo et al., 2003; Oertel and Young, 2004; Haenggeli et al., 2005; Zhan and Ryugo, 2006, 2007; Zeng et al., 2011; Balmer and Trussell, 2021, 2022). The idea is that sound source segregation is accomplished by standard binaural time and intensity data, pinna-based spectral cues, information about head and neck position with respect to gravity, and muscle proprioception for movement feedback that converge in the GCD. These multisensory lines, thereby making it a crucial structure of interest. Our interest was to investigate the nature of the multisensory inputs and to identify the cellular targets that help create three-dimensional auditory space.

The DCN is a cortical structure containing a rich variety of cell types and is situated dorsal to the posterior part of the ventral cochlear nucleus (Oertel & Young, 2004). The two principal divisions are separated by a thin but distinct layer of small cells composing a region called the granule cell domain (GCD, Mugnaini et al., 1980).

The granule cell domain (GCD) is not a homogeneous structure. It is characterized by granule cells (Mugnaini et al., 1980a), Golgi cells (Mugnaini et al., 1980b; Mugnaini, 1985; Ferragamo et al., 1998; Irie et al., 2006), two types of unipolar brush cells (Dino et al., 1994; Weedman et al., 1996; Borges-Merjane and Trussell, 2015), and Chestnut cells (Weedman et al., 1996). The inputs from auditory, somatosensory, vestibular, and brain stem reticular nuclei arrive as mossy fibers or boutons (Weedman and Ryugo, 1996; Wright et al., 1996; Ohlrogge et al., 2001; Ryugo et al., 2003; Haenggeli et al., 2005; Irie et al., 2006; Zhan and Ryugo, 2006, 2007). Inputs generate excitatory postsynaptic potentials (EPSPs) in neurons that exhibit high affinity uptake of glutamate and aspartate or strongly inhibitory outward potassium currents ((Haenggeli, Pongstaporn, Doucet, & Ryugo, 2005; Zeng, Shroff, & Shore, 2011).

The main output of the DCN is conveyed by pyramidal cells whose somata reside in layer II. Auditory nerve projections carrying information about sound terminate on the basal dendrites of pyramidal cells (Oertel & Young, 2004). In contrast, parallel fibers originating from the multimodal granule cells terminate on the apical dendrites of pyramidal cells (Wouterlood & Mugnaini, 1984; Wouterlood, Mugnaini, Osen, & Dahl, 1984). The parallel fibers, run along the long axis of the DCN, perpendicular to the isofrequency contours (Oertel & Young, 2004). The granule cells not only receive information about gravity, head, neck and pinna position, and body movement but their output is modified by local circuit excitatory and inhibitory input from Golgi cells, unipolar brush cells and chestnut cells. The anatomical nature of these connections will organize the circuits and determine the output to pyramidal cells.

It is proposed that the pattern of spike discharges expressed by these principal projection neurons is determined by their intrinsic properties and converging synaptic inputs from parallel fibers and auditory nerve fibers. Of particular interest is the mossy fiber. These endings are among the largest in the central nervous system (Ramón y Cajal, 1909; Lorente de Nó, 1981) and are expected to have strong drive on their target cells (Eccles et al., 1967; D’Angelo 2018). The strong drive, however, has shown considerable variation with respect to their target cells and coactivation through other microcircuits (De Zeeuw et al., 2011; Cosellato et al., 2015). Here, we seek to reveal details about the anatomical nature of these mossy fiber inputs to the GCD using serial section electron microscopy and three-dimensional model constructions. We have a special focus on the interactions among the C2 DRG, the cuneate nucleus (CuN), the lateral reticular nucleus (LRN), and the spinal trigeminal nucleus (Sp5), structures known to influence the type and quality of sound arriving at the ear.

## MATERIALS AND METHODS

This report is based on anatomical data collected from adult male, Sprague-Dawley rats (250-350 gm) acquired from Charles River. Animal care and surgery procedures strictly adhered to the guidelines of the NIH and were approved by the Johns Hopkins University Animal Care and Use Committees. For all anterograde tracing experiments, animals were deeply anesthetized by an intraperitoneal injection of sodium pentobarbital (45 mg/kg body weight). Oral secretions were blocked by 0.05 mg intramuscular injections of atropine sulfate. Surgery was performed only after animals were unresponsive to paw pinches.

### Injection procedure and histological processing

Anesthetized animals were positioned into a stereotaxic frame, the skin and soft tissue over the parietal bone were reflected, and a 2-mm-diameter hole was drilled in the skull. A glass pipette (30-50 µm ID), filled with a mixture of 2.5% biotinylated Phaseolus vulgaris leucoagglutinin (PHA-L; Vector Laboratories, Burlingame, CA) and 2.5% PHA-L (Vector Laboratories) in phosphate buffer, pH 7.9, was lowered into the brain using a micromanipulator and an oocyte injector (NanoInjector, Drummond Instruments). The pipette tip was positioned according to stereotaxic coordinates so that its tip was placed in the left cuneate nucleus, spinal trigeminal nucleus, or lateral reticular nucleus guided by a rat stereotaxic atlas (Paxinos and Watson, 1982). For the dorsal root ganglion, the cervical vertebrae were exposed after a midline incision and dissection of the neck muscles. The first vertebra was partially removed to expose the C2 DRG and spinal nerve. Multiple injections of the tracer using a total volume of 0.5-1 µL were made into the ganglion. When all injections were completed, the exposed muscle and skin were cleaned, and the skin was sutured. The animal was allowed to recover in its home cage.

After 10-14 day post-injection survival times, rats were anaesthetized with a lethal dose of pentobarbital and transcardially perfused with chilled 0.12 M phosphate-buffered (PB) 4% paraformaldehyde (pH 7.3). The brains were removed and kept in the same fixative for an hour, embedded in 5% gelatin-albumin hardened by paraformaldehyde, and then sectioned in the transverse plane using a vibrating microtome. Consecutive sections (50 µm thick) were serially collected in culture wells containing 0.12 M PB. The sections were then treated with biotinylated peroxidase-avidin complex (ABC-*Elite^TM^*, Vector, Burlingame, CA) and Ni-DAB to produce an electron-dense reaction product that was visually detectable by light and electron microscopy.

The sections containing labeled endings for analysis using the electron microscope were flat-embedded in PolyBed 812 (Polysciences, Inc.,Warrington, PA) between two sheets of Aclar (Electron Microscopy Sciences, Hatfield, PA).

### Transmission Electron Microscopy

EM procedures standardized by this lab have been described previously (Haenggeli et al., 2005; Wright & Ryugo, 1996; Zhan, Pongstaporn, & Ryugo, 2006; Zhan & Ryugo, 2007). Generally, the selected sections with positive labeling in the GCD were osmicated in 1% OsO_4_, rinsed thoroughly in buffer, stained in 1% uranyl acetate for 1 hr, rinsed again repeatedly, and finally dehydrated in ascending concentrations of ethanol. The tissue that was flat embedded in Epon were analyzed by light microscopy and maps of selected regions were made for orientation purposes, using blood vessels and the labeled structures as landmarks. We dissected specific regions containing labeled endings from these tissue sections and re-embedded the smaller pieces in BEEM capsules for ultrathin sectioning. Serial ultrathin sections were cut at 75 nm, mounted on Formvar-coated slotted grids, and examined and photographed using a Hitachi H-7600 electron microscope.

### Data analysis

Electron micrographs were collected from identified mossy fiber and bouton terminals in the DCN that arose from a successful injection confined to the cuneate nucleus (n = 5), spinal trigeminal nucleus (n = 4), lateral reticular nucleus (n = 6), or the C2 DRG (n = 7). The anterogradely labeled mossy fiber endings were reliably distributed in the granule cell domain. Each of the analyzed endings contained from 7 to 101 serial sections (Table 1), representing a bouton or mossy fiber. The photos of selected structures were aligned in register to each other by rotation and translation until corresponding structures of adjacent sections were superimposed. Endings including mitochondria and postsynaptic densities were manually traced so that three-dimensional (3D) surface reconstructions could be generated and volumes calculated by *Amira 4.1.* (Mercury Computer Systems, Berlin). For synaptic vesicle analysis, the electron micrographs were initially imported into photoshop, and the vesicles were highlighted by a “blinded” researcher. The highlighted photographs were then transferred to ImageJ. Both the long and short axis were automatically determined by *ImageJ 1.54a*. The data were further transferred to *GraphPad* (Boston, MA) for visualization and *Matlab2022a* (Natick, MA) for statistical analysis.

**Table 1.**
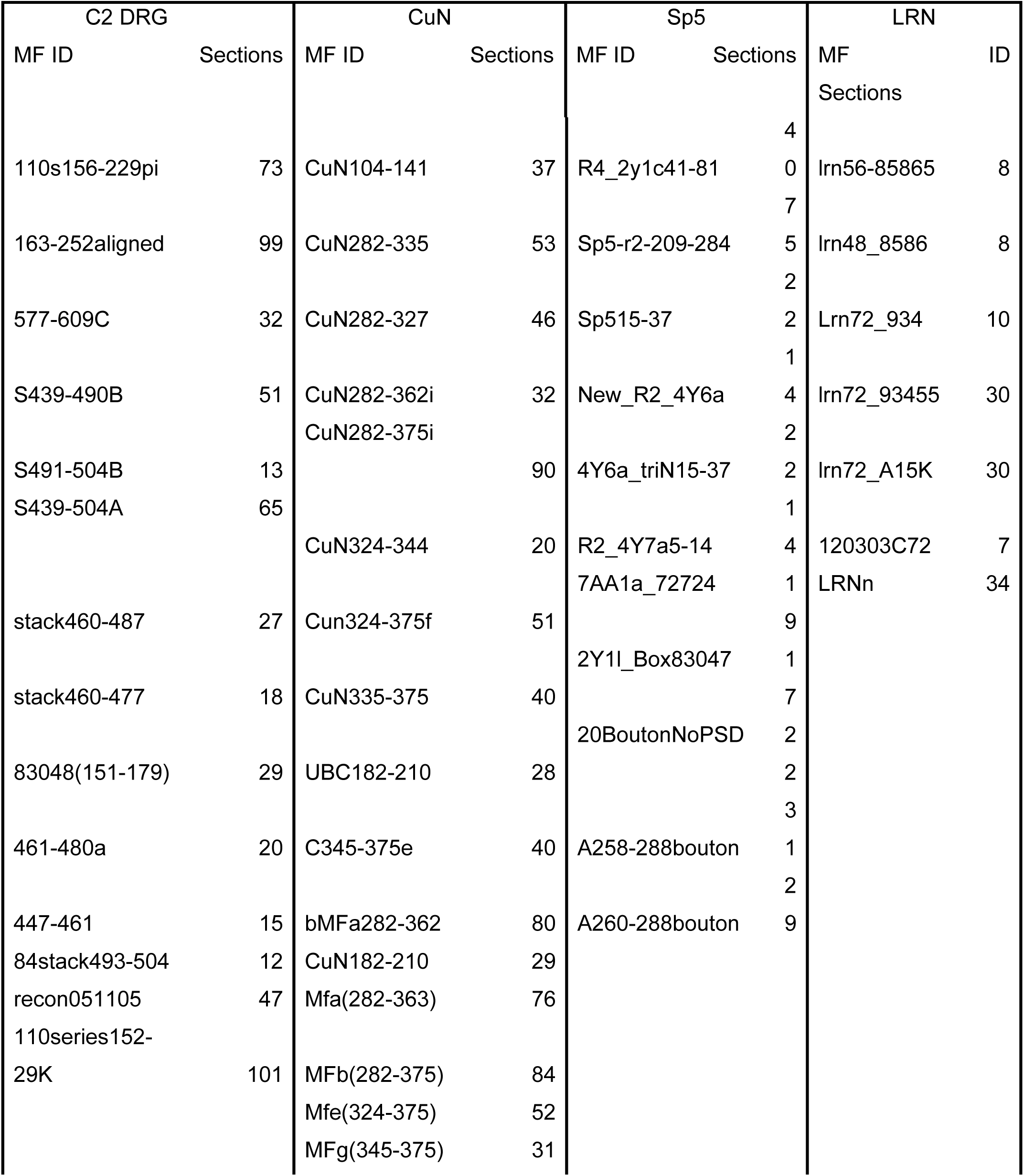
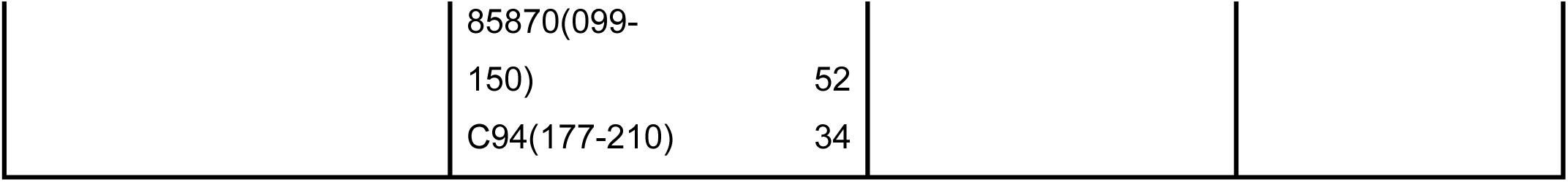
Mossy fiber or bouton endings from MSN projections.

## RESULTS

Mossy fibers are terminal endings that are large and rather complex in shape, ranging from slightly disfigured spheres to elongated, elaborately twisted, convoluted forms with irregular surfaces and varying girth. Mossy fibers can be up to 20 µm in length with exhibit irregular shapes as seen under brightfield light microscopy (**Figure 1A**), but the detailed structure can best be appreciated using confocal microscopy with multidimensional digital analyzers (Balmer and Trussell, 2019) or electron microscopy (**Figure 1B**). The mossy fiber is highly irregular with hair-like extensions and is complemented by the postsynaptic target which can also emit penetrating by filiform hairs. The mossy fibers examined in this report were localized to the region called the GCD of the cochlear nucleus, which appears as a thin microneuronal shell residing over the medial, dorsal, and lateral surface of the ventral cochlear nucleus and extends into layer II of the dorsal cochlear nucleus as described by Mugnaini et al. (1980a). Most of our analyses was performed in the lamina, which is the sheet of microneurons separating the dorsal cochlear nucleus above from ventral cochlear nucleus below, and to a lesser extent in superficial granule cell layer overlying the dorsolateral surface of the ventral cochlear nucleus and the subpeduncular corner, located at the dorsomedial edge of the ventral cochlear nucleus just below the cerebellar peduncles.

**Figure 1.**
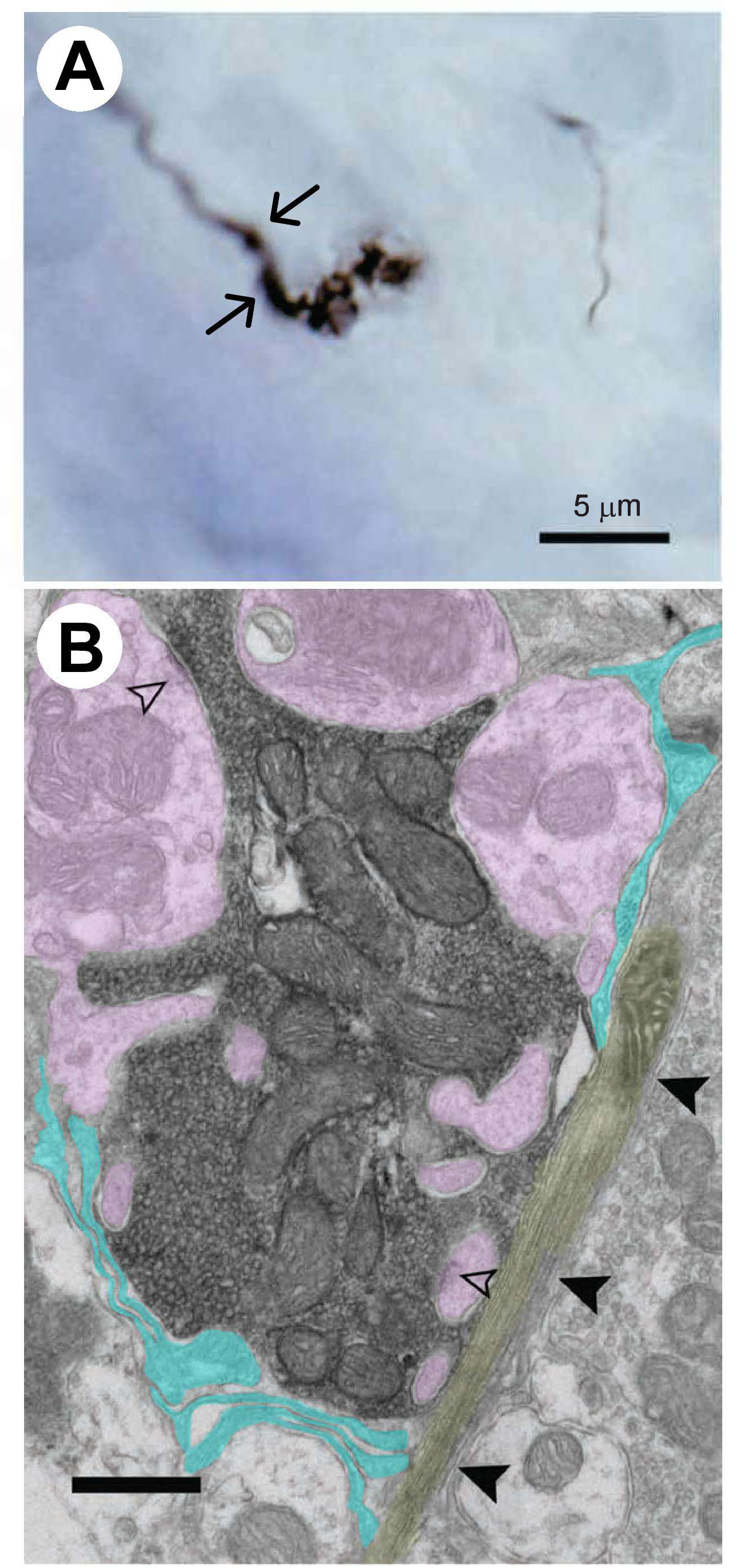
Mossy fibers are revealed in light and electron microcopy. **(A)** Light micrograph shows a mossy fiber from the cuneate nucleus in the lamina of the GCD in the cochlear nucleus. After two *en passant* swellings (arrows), this mossy fiber emerges from the end of an axon as two prominent and irregular varicosities. **(B)** An electron micrograph of a mossy fiber from the cuneate nucleus. It is nestled into the claw of a granule cell dendrite and penetrated by filiform dendrioles (pink coloring). The glomerular complex is incompletely encased by glial lamella (light blue) and a thin unmyelinated axon (highlighted in yellow, arrowheads) of unknown origin. Synapses on dendrites are indicated by open arrowheads. Scale bar: A, 5 µm; B, 0.5 µm.

The receptive targets of the mossy fiber are equally complex in spatial terms. The granule cell can give rise to several dendrites, each of which terminates as an elaborate claw with branches that are postsynaptic to the incoming mossy fiber. Filiform dendritic extensions (approximately 120-150 nm in thickness) can emerge from the claw and to penetrate the mossy fiber, serving as anchors for adhesion. Sometimes, multiple mossy fibers can contact a single granule cell claw. Other times, dendrites from separate granule cells and UBCs can contact the same mossy fiber. This complex of mossy fiber and recipient dendrites is joined by axon terminals with round or pleomorphic synaptic vesicles and encased by a discontinuous glial sheath. The structure is called a glomerulus (Jones and Powell, 1969; Peters et al., 1976; Ribeiro-da-Silva and Coimbra, 1982; Ribeiro-da-Silva et al., 1985; Mugnaini et al., 2011).

### The microcircuitry of mossy fibers Cervical Dorsal Root Ganglion

We examined the postsynaptic targets of 14 mossy fibers that were labeled by unilateral injections of BDA into the dorsal root ganglion (DRG) of the second cervical spinal root. These mossy fibers terminated in a GCD glomerular claw (n = 9) that included axons (n = 4) of unknown sources or within the tuft of UBC dendrites (n = 4) (**Figure 2**). The size of UBC dendritic shafts makes it an accessible postsynaptic target, and large PSD are frequently encountered along the apical shaft and terminal dendrites of UBCs (**Figure 3 and 4**). The PSDs of granule cells are small by comparison. In our material, C2 DRG mossy fibers produced 3 - 10 PSDs across a length of 2 - 5.8 µm in length. In addition, mossy fibers were occasionally observed to branch and give rise to bouton endings as well as to be postsynaptic to inputs of unknown origins (**Figure 2A**).

**Figure 2.**
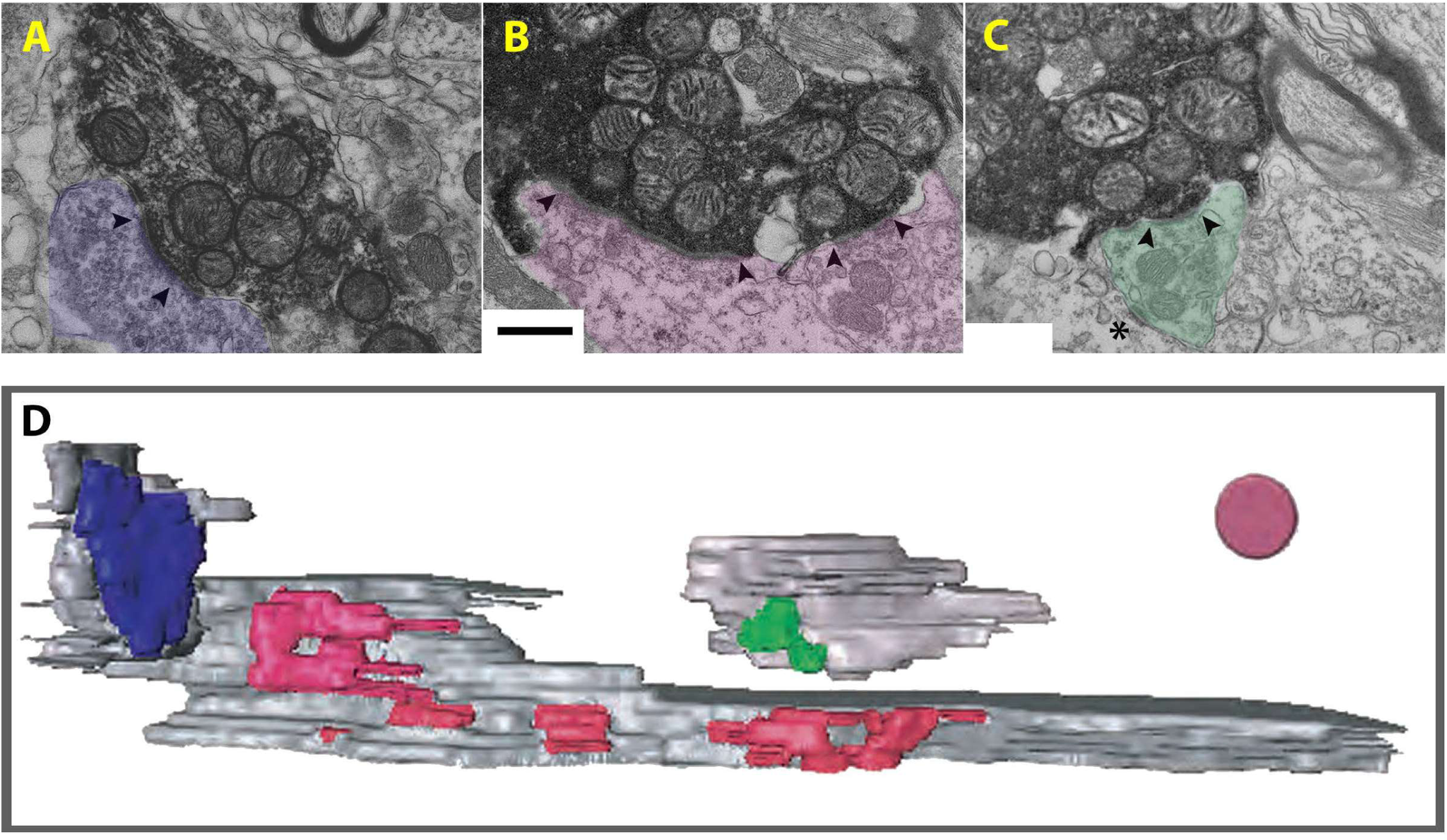
Electron micrograph of a DRG mossy fiber and its major targets. **A, B, C**: Labelled C2 DRG mossy fiber endings are indicated in the EM photos (upper panel, black reaction product). **(A)** A mossy fiber forms a synapse (arrowheads) with an axon of unknown origin (blue). **(B)** This mossy fiber forms a long and a short synapse against a large dendrite. The long synapse is characteristic for a UBC. **(C)** A labelled mossy fiber synapses with a dendritic spine (marked by arrowheads) of presumable stellate neuron, that in turn forms a gap junction with dendritic spine (**\***) of unknown origin. **(D)** 3D reconstruction shows the postsynaptic densities of the C2 DRG axonal terminals and postsynaptic targets. The mossy fiber (hidden for clarity) synapses against an axon terminal (blue), forms several synapses (pink) with a large UBC dendritic shaft, and synapses with the dendritic spine of a granule cell (green). Scale bar and disc diameter = 0.5 µm.

**Figure 3.**
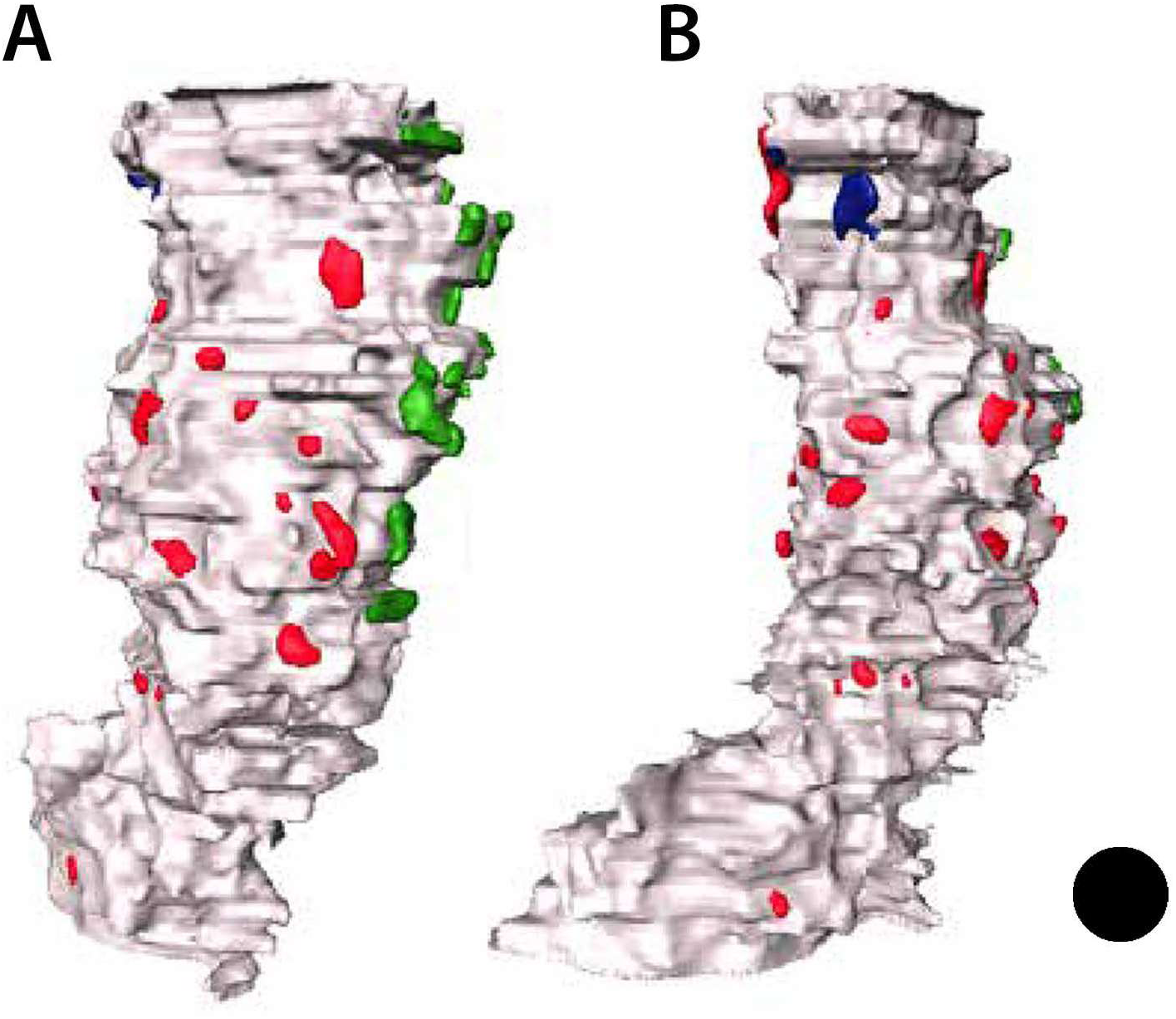
3D computer reconstruction of a UBC dendritic shaft. The image of the dendrite on the left (A) has been rotated 180 degrees from the image on the right (B). Note the irregular surface of the dendrite and the relatively nonuniform distribution of PSDs (red) from the C2 DRG mossy fiber. All PSDs on this dendrite was reconstructed. The source of the other PSDs (blue and green) is unknown. Scale bar equals 1.0 µm.

**Figure 4.**
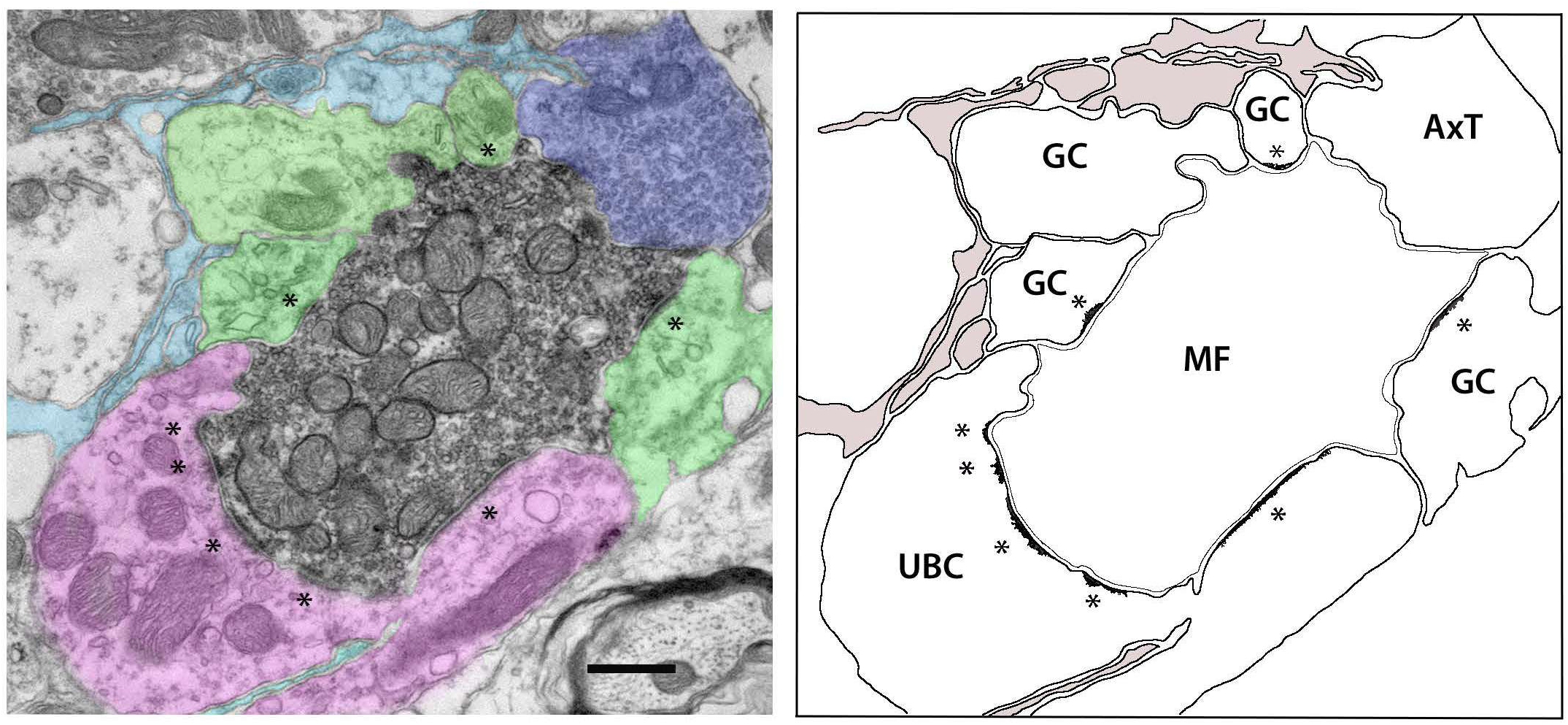
A mossy fiber from C2 DRG reveals multiple postsynaptic targets. **A.** The electron micrograph shows that postsynaptic targets are UBC dendritic shafts (pink), granule cell dendrites (green), and an axon terminal (blue) with pleomorphic synaptic vesicles from an unknown source. Part of this glomerulus is lined by glial sheaths (light blue). Synapses are indicated by asterisks. **B.** Outline of the elements of this glomerulus with labels. Glia is colored light brown. AxT, axon terminal of unknown origin; GC, granule cell dendrite; MF, mossy fiber from the second cervical dorsal root ganglion; UBC, unipolar brush cell dendrite. Scale bar equals 0.5 µm.

The UBCs have a single, thick dendritic stalk whose terminal dendritic tuft resembles the apical tentacles of a sea anemone. The tuft can completely engulf a single mossy fiber (see Figure 4 of Weedman and Ryugo, 1996). UBCs have few or no synaptic inputs onto their cell body nor on the basal stretch of their single dendritic shaft (Dino & Mugnaini, 2000; Floris, Dino, Jacobowitz, & Mugnaini, 1994; Mugnaini, Dino, & Jaarsma, 1997; Mugnaini & Floris, 1994). At the apical end of the UBC dendritic shaft, however, the thin dendritic branches mark the start of synaptic inputs. Granule cell dendrites are also involved in this plexus and occasionally receive input from an unlabeled axon.

### Cuneate Nucleus

The cuneate mossy fibers are diverse in size, where some of the larger ones are penetrated by their postsynaptic targets. The mossy fiber can undulate in diameter, forming multiple swellings that can be as wide as 7 µm across with a single swelling forming as many as forty-six synapses. This characteristic feature of MFs substantially increases the contact surface between the ending and its targets, which may make synaptic transmission more efficient in passing along information. The terminal dendritic tufts of UBCs and the dendritic claws of granule cells also enlarge the target space for mossy fibers.

On the basis of analyzing mostly uninterrupted serial strings of EM sections involving another labelled mossy fiber, we could infer structure continuity with confidence and determined that 12 postsynaptic targets of mossy fibers were dendrites of granule cells and six were dendrites of UBCs (**Figure 5**). Three additional mossy fibers were part of this glomerulus that targeted granule cell dendrites, which were also postsynaptic to axons of unknown origin. Another cuneate fiber appeared to have two UBC dendrites (green) are the postsynaptic targets of this cuneate mossy fiber (Figure 6). The dendrites are identifiable by their content of free ribosomes and long asymmetric PSDs. A branch of the cuneate mossy fiber terminates on the opposite side of one UBC dendrite. In addition, an axon terminal of unknown origin but containing round synaptic vesicles makes synapses with this UBC dendrite (Yellow). The 3D reconstruction of this synaptic site shows the interleaving of UBC dendrites and mossy fiber as well as a contribution from another terminal (Figure 6B).

**Figure 5.**
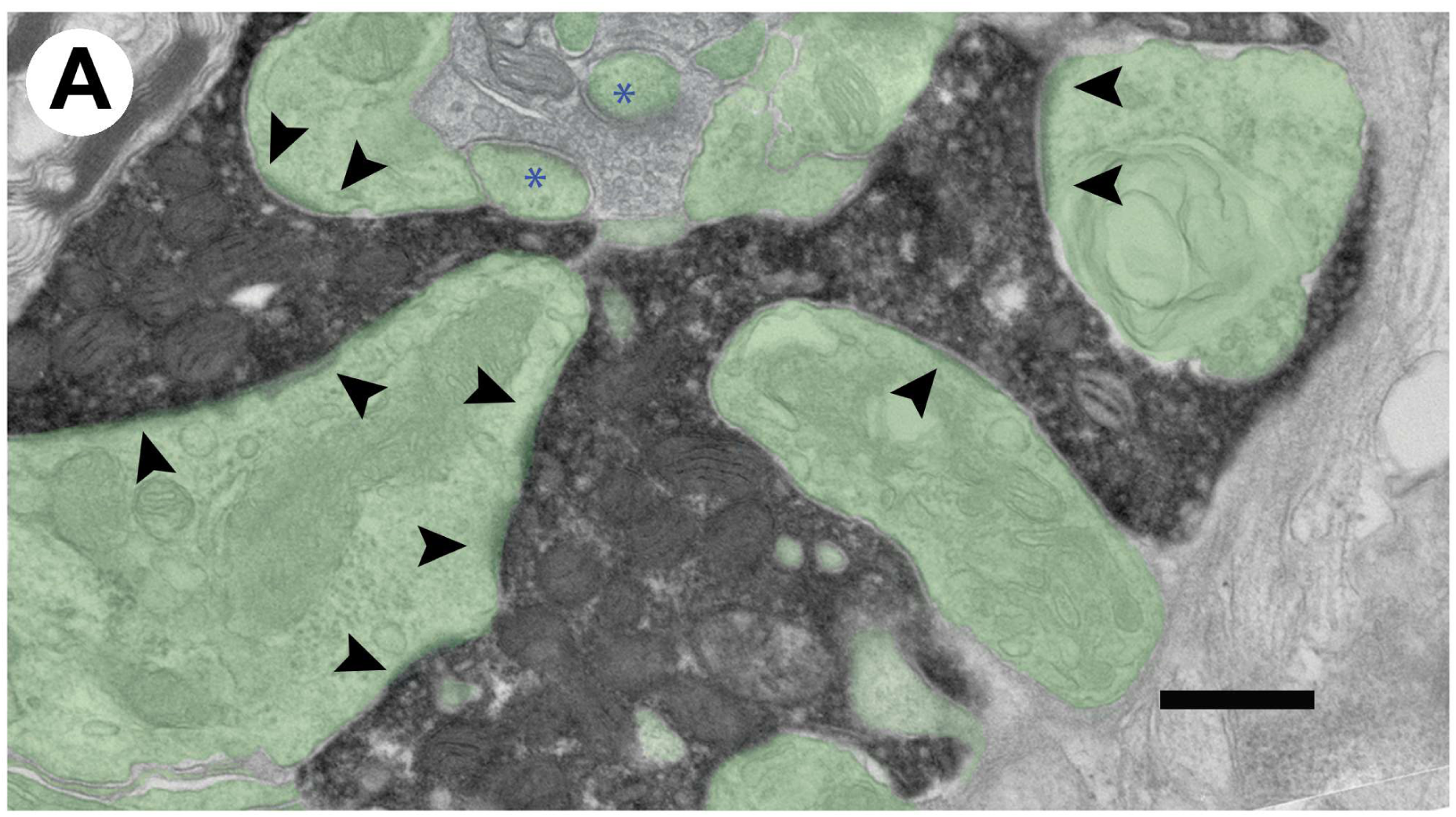
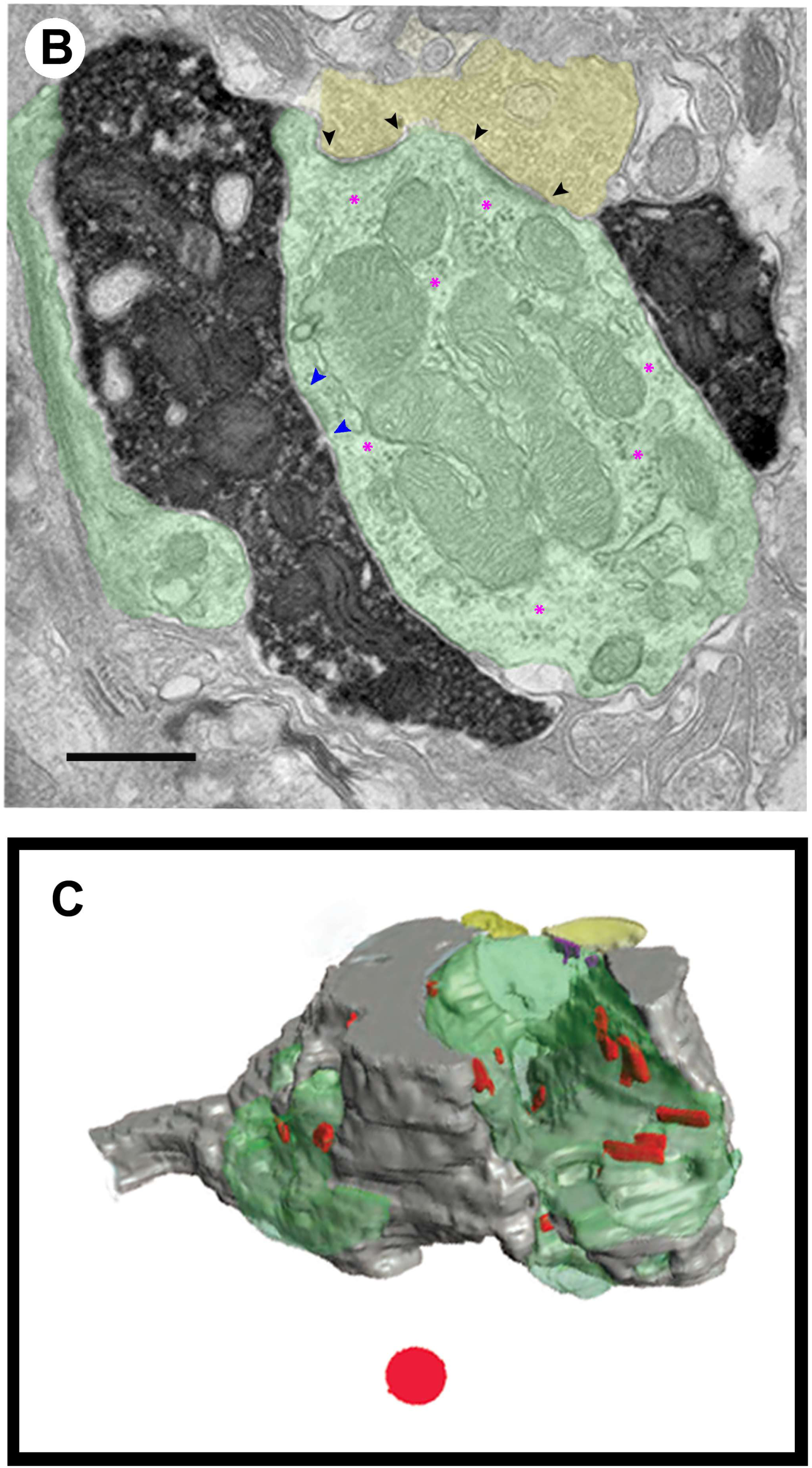
Electron micrograph shows a labelled mossy fiber from the cuneate nucleus that forms synapses (black arrowheads) with dendrites from the apical tuft of a UBC (green, A). An unlabeled axon terminal is part of this glomerulus and forms synapses (blue asterisks) with thin dendrites assumed to be part of the UBC dendritic tuft. Both terminals contain round synaptic vesicles. Serial section analysis revealed that these postsynaptic targets arose from a single UBC. The source of the thin dendrites embedded in the unlabeled mossy fiber could not be identified. **(B)** The labelled mossy fiber (black precipitate, 750 nm from **A**) is flanked by 2 UBC dendrites (green) and forms a synapse with one (blue arrowheads). Free ribosomes (magenta asterisks) are scattered around the cytoplasm, identifying the dendrite as belonging to a UBCs. An unlabeled terminal (yellow) containing round synaptic vesicles forms 2 asymmetric synapses with the UBC (black arrowheads). (**C)** This 3D reconstruction is based on serial sections of the axon shown in panel **A.** The axon enters from the left. The surface of the mossy fiber (gray) is revealed with its irregular shape, the flanking UBC dendrites (green), and the branching axon terminal (yellow). PSDs are shown in red. Scale bar and circle equal 0.5 µm.

**Figure 6.**
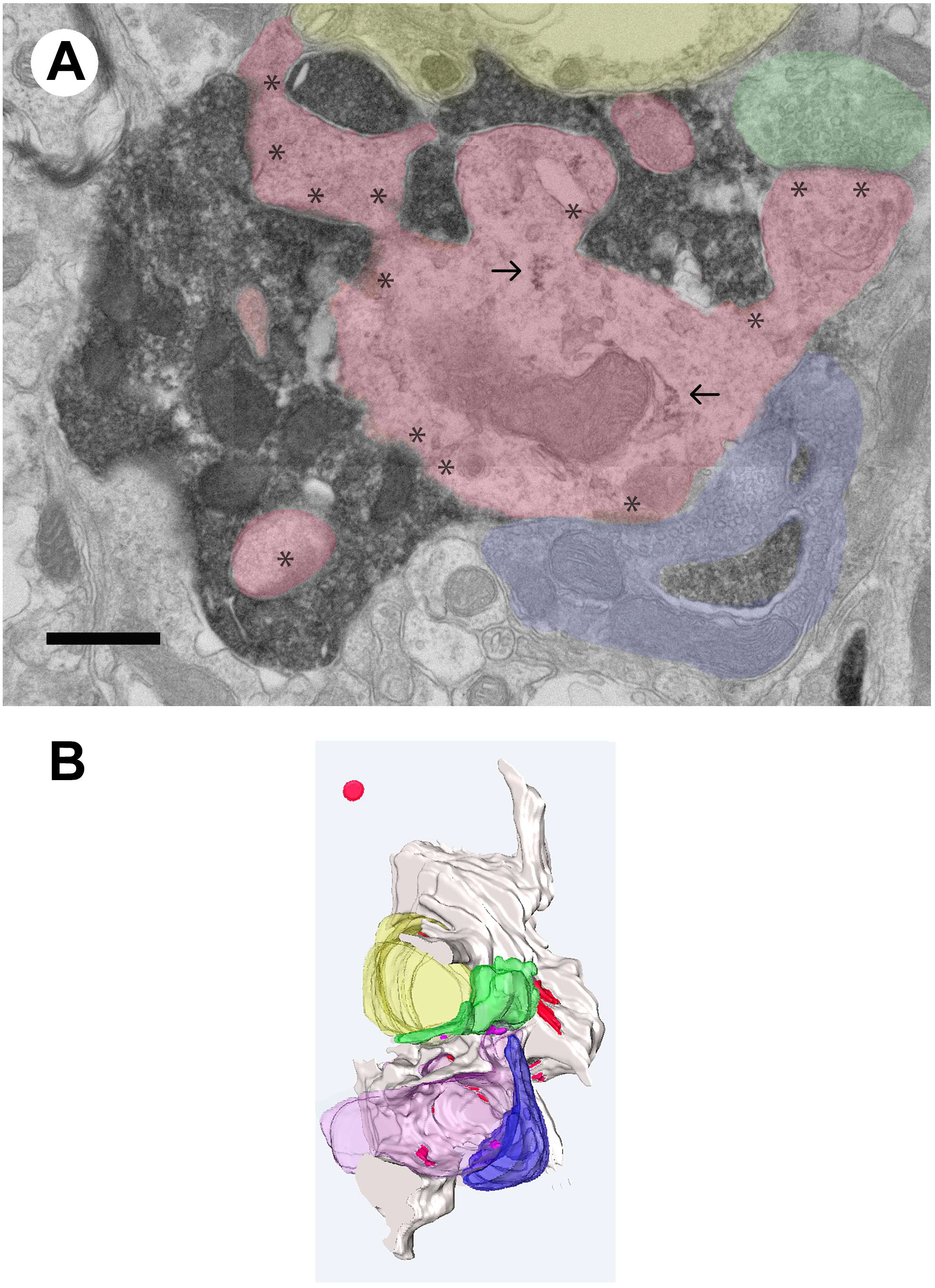
A labelled mossy fiber from the cuneate nucleus that terminates in the GCD. **(A)** Electron micrograph shows the labelled mossy fiber (black precipitate) with round synaptic vesicles making multiple synaptic contacts (*) within a glomerulus formed by the apical tuft of UBC dendrites (pink). Free ribosomes that characterize UBC dendrites are indicated by arrows. This dendrite is also postsynaptic to 2 axon terminals of unknown origin (green and blue). Both of these additional terminals contain round synaptic vesicles. A branch of the labelled mossy fiber penetrates the blue axon in 2 places. This axon terminal (blue) is also presynaptic to the main dendrites of the glomerulus (pink). A third terminal with round synaptic vesicles (green) also synapses on the main glomerular dendrite. **(B)** 3D computer rendering of the glomerulus with the labelled mossy fiber (light gray) with a UBC (pink) and a granule cell (yellow) dendrite plus two bouton endings (blue and green). Synapses are indicated in red. The neuropil is obviously quite complex. Scale bar and circle equal 0.5 µm.

The interleaving of a mossy fiber with elements of the postsynaptic target is illustrated by the relationship with this LRN mossy fiber and the dendritic tuft at the apex of a UBC dendrite (**Figure 6**). Further evidence for the UBC identification is the presence of ribosomes in the cytoplasm (arrows). Note how the mossy fiber nestles in among the multiple branches of the tuft and forms a number of asymmetric synapses (asterisks). This glomerulus has other components of undetermined origins including 2 axon terminals containing round synaptic vesicles and a dendrite that receives synapses from the labelled terminal in sections out of this plane of view.

### Lateral Reticular Nucleus

The electron microscopic examination of a mossy fiber originating from an axon of the lateral reticular nucleus was shown to form multiple synapses (**Figure 7A, B**). The 3D reconstruction of this swelling revealed a volume of 2.01 µm^3^ and surface area of 8.23 µm^2^. The mitochondria occupy a volume of 0.267 µm^3^, which is 13.2% of the calculated total bouton volume. The ratio of bouton volume to surface area is 0.244 µm. This ending exhibits the typical core of mitochondria and round synaptic vesicles opposing asymmetric membrane thickenings. Such endings are presumed to be excitatory.

**Figure 7.**
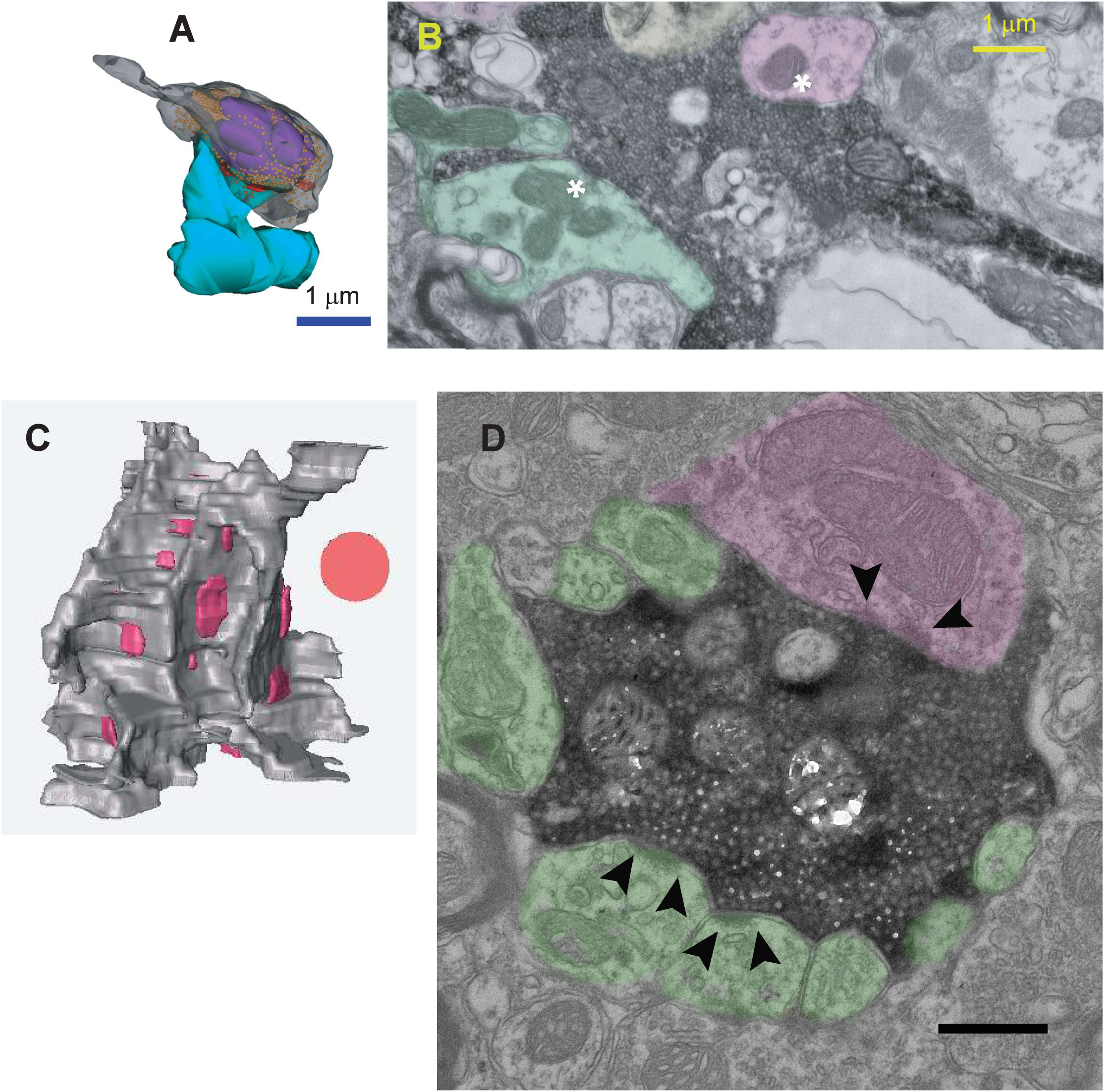
Mossy fiber bouton (gray) from the lateral reticular nucleus, is exhibited in 3D reconstruction in the GCD. 3D rendering of the bouton (transparent for clarity) that makes two synapses with dendritic spines (blue). The bouton has a typical core of mitochondria (purple) with scattered synaptic vesicles (orange). **B.** An electron micrograph shows the synapses, and their targets highlighted in color and asterisks. **C.** 3D reconstruction of another LRN mossy fiber indicating multiple PSDs (pink). **D.** An electron micrograph shows the labelled mossy fiber forming synaptic contacts with dendrites of granule cells (green) and a unipolar brush cell (pink). C, D. Scale bar and circle equal 1 µm.

Another LRN mossy fiber displays more complicated structure and more postsynaptic targets as appeared originated from other sources (Figure 7C, D). One UBC dendrite (pink) is a postsynaptic target of this LRN mossy fiber, whereas the remaining postsynaptic elements appear to be granule cell dendrites (green). The dendrites are identifiable by the presence of free ribosomes and elongated asymmetric postsynaptic densities (PSDs). Three-dimensional reconstruction of this synaptic site reveals a complex architecture with multiple protrusions that segregate distinct PSD-associated release sites. There were labelled endings from the LRN containing pleomorphic vesicles observed with an associated symmetric PSD (**Figure 8**). Such endings were rare, presumed to be inhibitory, and found among other endings containing round synaptic vesicles. Because we did not stain for “inhibitory” neurons as part of this investigation, we cannot comment as to their chemical nature. The dendrite contacted by this LRN ending containing pleomorphic synaptic vesicles was also targeted by terminals filled with round synaptic vesicles. We see two mossy fiber endings, one with round synaptic vesicles and one with pleomorphic synaptic vesicles interacting with two dendrites. Both dendrites receive synapses from a presumed excitatory mossy fiber because of its round synaptic vesicles, whereas one dendrite receives input from what appears to be an inhibitory mossy fiber.

**Figure 8.**
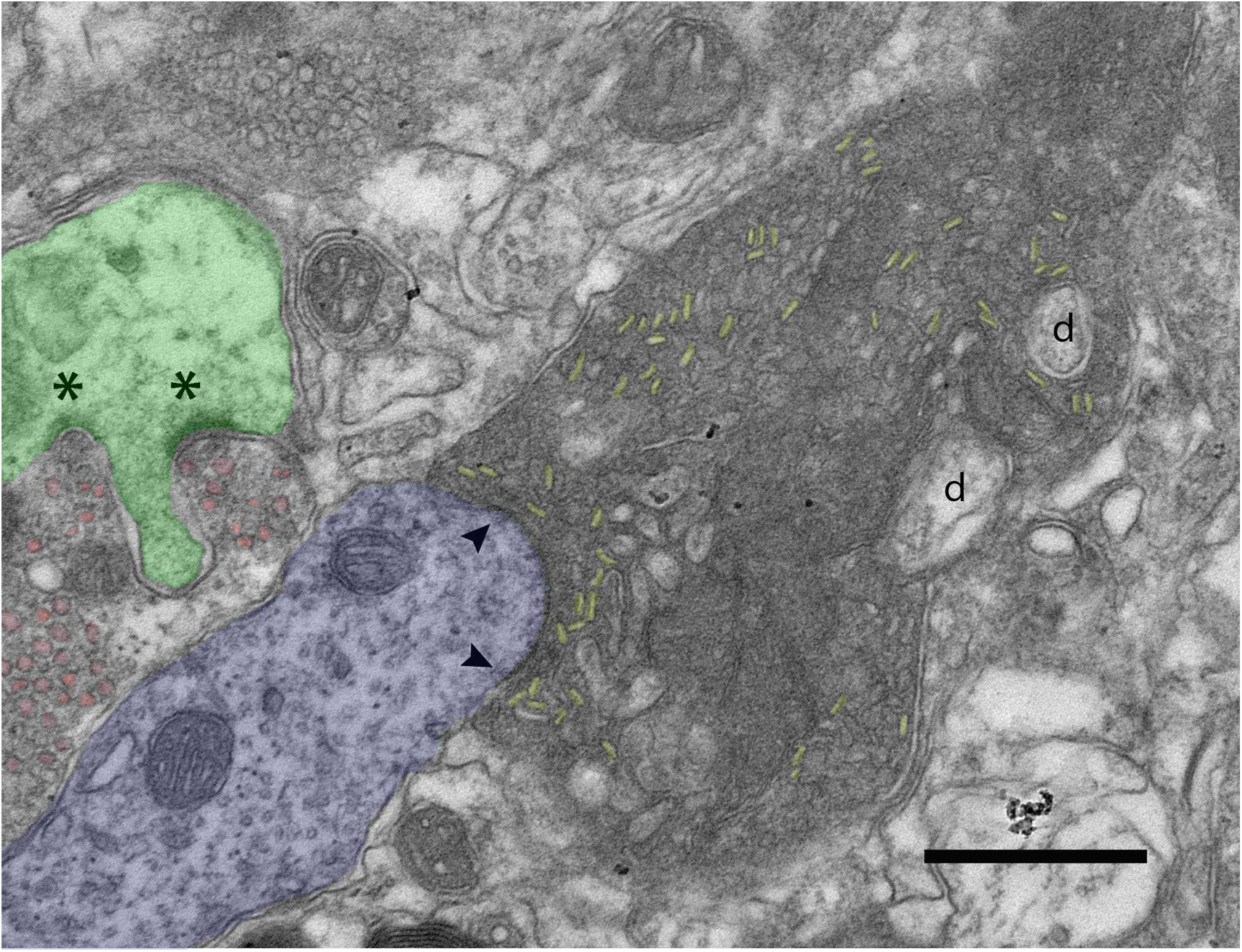
A bouton from the lateral reticular nucleus forms a synapse (arrowheads) with a dendritic shaft (blue). The dendritic shaft displays a long PSD as well as the presence of free ribosomes, features characteristic of UBC dendrites. This labelled ending contains pleomorphic synaptic vesicles, which is consistent with the symmetric PSD. The postsynaptic dendrite also receives a synapse (not in this plane of section) from the unlabeled ending with round synaptic vesicles (pink) that also synapses (**\***) on the adjacent dendrite (green). A third ending in this micrograph contains round synaptic vesicles but appears isolated from the green dendrite by glial laminae. Scale bar equals 0.5 µm.

### Spinal trigeminal nucleus

Granule cells of the GCD are targets of spinal trigeminal projections as previously reported (Haenggeli et al., 2005). Electron micrographs of Sp5 boutons reveal a terminal filled with round synaptic vesicles and exhibiting a somewhat scalloped perimeter (**Figure 9**). This bouton has a volume of 3.496 µm^3^, and a surface size of 17.71 µm^2^. The mitochondria of this bouton represent about 24.1% of the volume fraction, which is similar to that of LRN boutons. This bouton has three PSDs in this plane of section but makes synapses with all dendrites at different levels above and below this plane.

**Figure 9.**
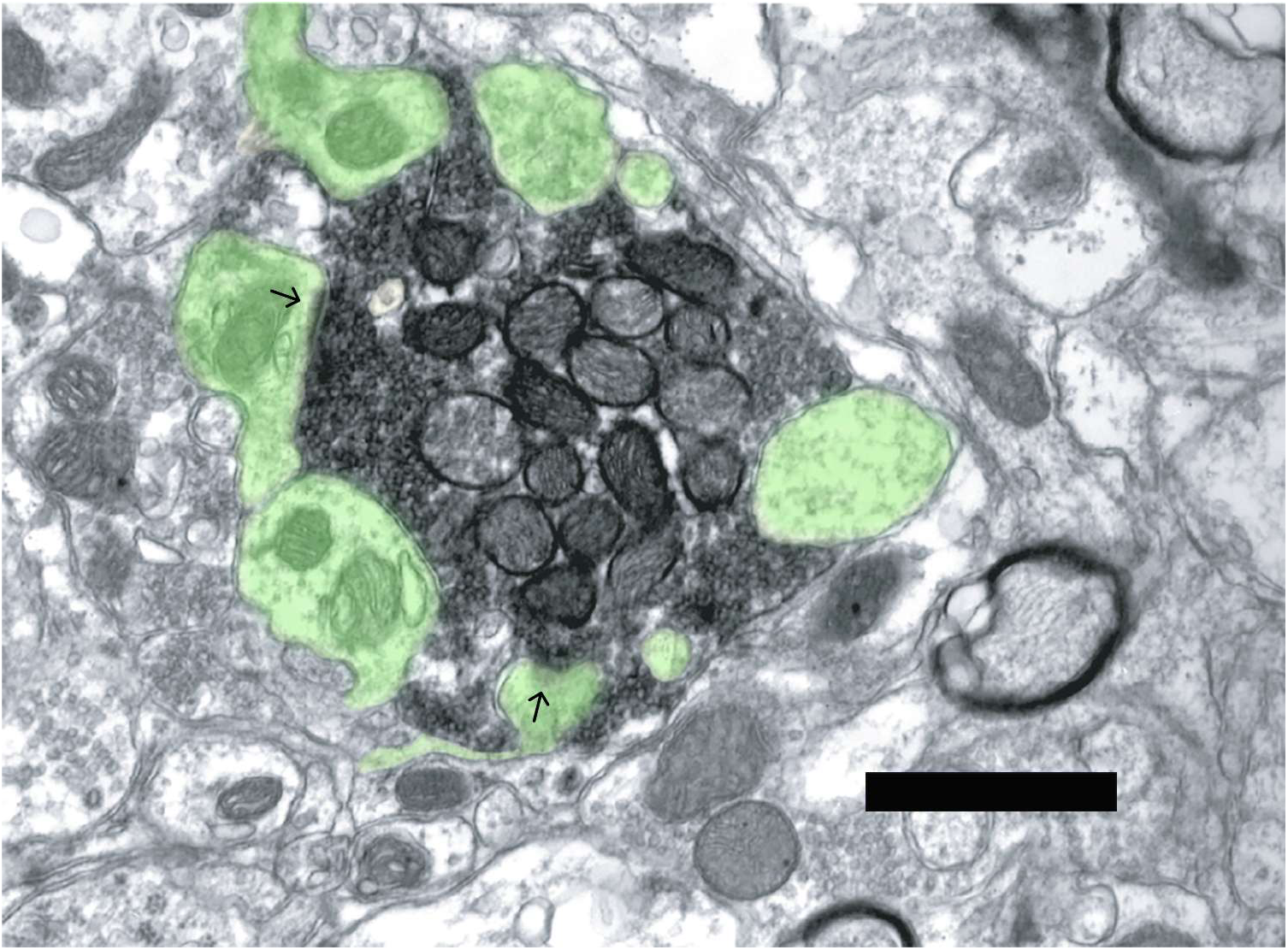
An electron micrograph shows a labelled bouton (black reaction product) from the spinal trigeminal nucleus that terminates in the granule cell domain. This bouton shows a central core of mitochondria and scattered round synaptic vesicles within the cytoplasm. The terminal surface is scalloped by indentations of postsynaptic dendrites (green). Serial sections showed that each dendritic profile in this glomerulus received at least one synapse (arrows) from this ending. Scale bar equals 0.5 µm.

Sp5 mossy fibers are found to oppose exceptionally large PSDs with thick dendrites containing free ribosomes, implying contact with UBCs (**Figure 10**). In this case, the mossy fiber containing round synaptic vesicles, penetrates well into the dendrite. The mossy fiber forms 2 long asymmetric PSDs, typical for UBCs. Our 3D reconstruction of the mossy fibers reveals that the synapses are fenestrated, a feature primarily found with very large synapses. The separate pieces of the dendrite are joined by a desmosome near the apex of the mossy fiber.

**Figure 10.**
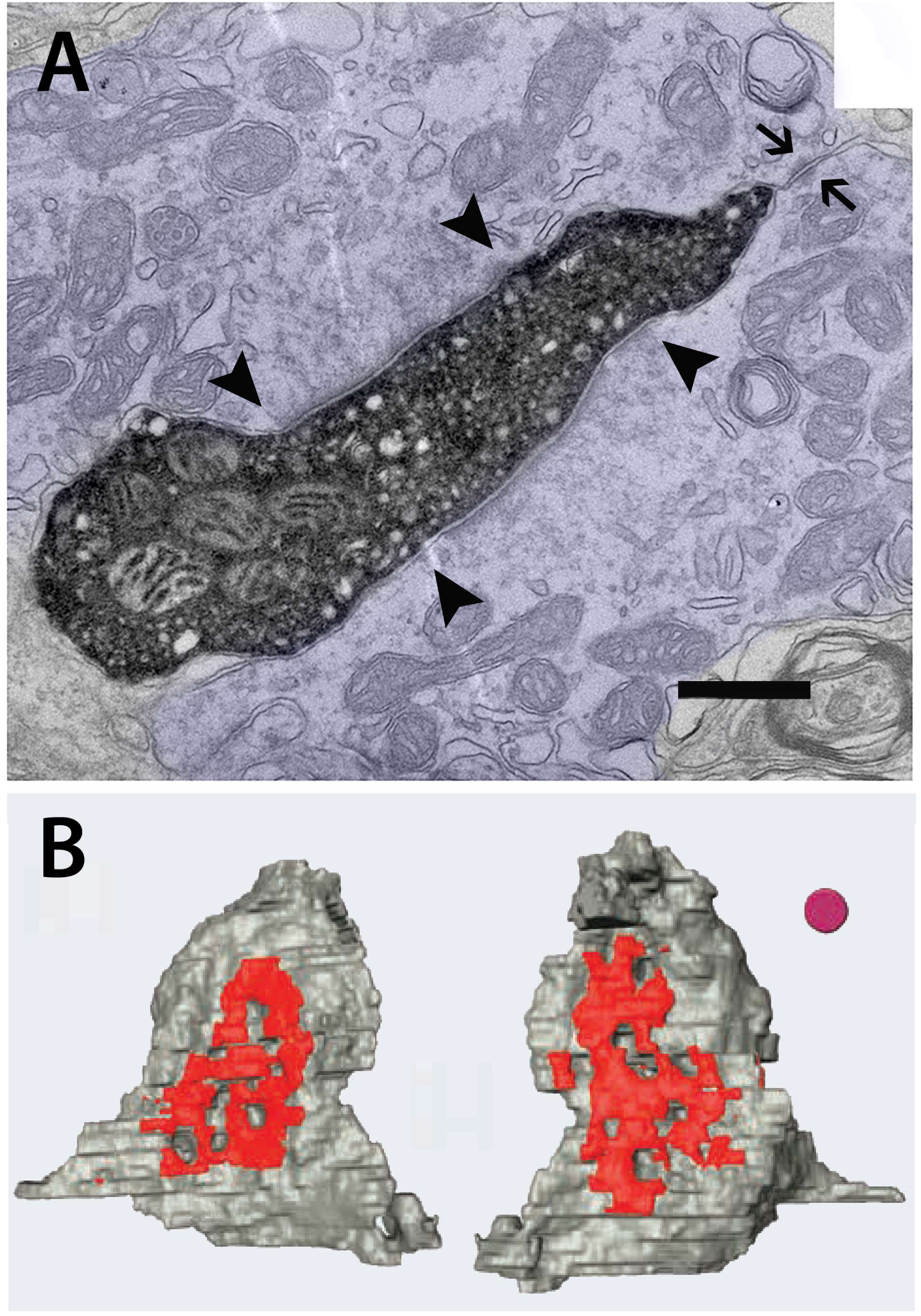
**(A)** This electron micrograph shows a labelled ending (black reaction product) from the spinal trigeminal nucleus inserted into a thick UCB dendrite. The mossy fibers with round synaptic vesicles is associated with two asymmetric postsynaptic densities (arrowheads) of significant length. Note also that the separate dendritic leaves are joined by a desmosome (arrows). The mossy fiber penetrates a large dendritic shaft with two groups of active zones (red) on opposite sides (upper and lower panels). **(B)** The 3D reconstruction of this mossy fiber reveals that it forms two large PSDs. Each PSD is among the largest in our mossy fiber population. The presence of fenestrations in the PSD serves to keep unbound transmitter in close proximity to binding and reuptake sites. Scale bar and disk equals 0.5 µm.

In addition, there are Sp5 endings targeting the soma of a likely Golgi cell (**Figure 11**). Golgi cells have prominent Golgi organization and rough endoplasmic reticulum in their cell bodies. This particular neuron has an eccentrically positioned nucleus and many cisternae scattered around the cytoplasm. The terminal is filled with pleomorphic vesicles and exhibits symmetric pre- and postsynaptic densities, suggestive of inhibitory action at the synapse. Such a relationship is inferred to have an overall excitatory effect because of its inhibitory action on the inhibitory neuron. Because Golgi cells provide inhibition to hundreds of granule cells, their silence can exert massive disinhibition on downstream granule cells (Yaeger & Trussell, 2015).

**Figure 11.**
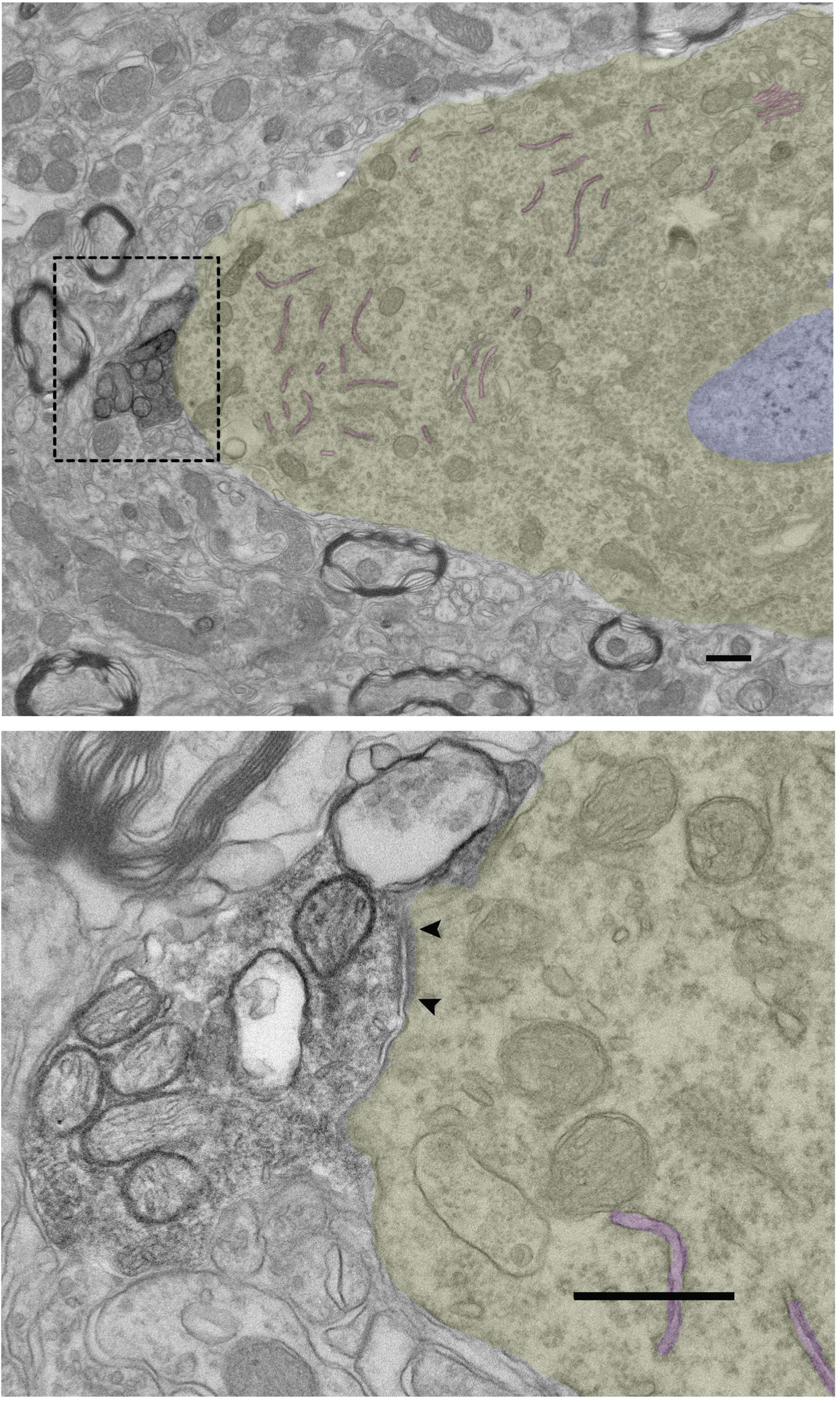
An electron micrograph of a labelled bouton (black reaction product) from the spinal trigeminal nucleus that targets a presumed Golgi cell (yellow coloring). **(A)** This postsynaptic cell was identified by its eccentric nucleus (blue), multiple stacks of Golgi cisterns, and numerous rough endoplasmic reticulum (pink). **(B)** An enlarged view of the labelled terminal taken 7 sections away from that shown in **A**. The terminal contains flattened synaptic vesicles that oppose a symmetric postsynaptic density (arrowheads). An inhibitory terminal that synapses on an inhibitory neuron implies a release from inhibition and a spread of excitation. Scale bar equals 0.5 µm.

### The synaptic vesicles of sensory mossy fibers

Although synaptic vesicles all appear round when rapidly frozen (Tatsuoka and Reese, 1989), when fixed by aldehyde solutions and examined under the electron microscope, vesicles at symmetric synapses appear more susceptive to flattening (Rizzoli & Betz, 2005). They may have different morphological properties in terms of their size, shape and postsynaptic targets, and the key question is whether vesicle shape indicates synaptic action. A mossy fiber typically has a large pool of round synaptic vesicles occupying a large volume of mossy fibers. The synaptic vesicles surround the mitochondria core of the mossy fibers or boutons (**Figures 1, 10**). There are, however, instances where a labelled mossy fiber contains pleomorphic or at least flattened vesicles (Figure 8). Terminals containing pleomorphic synaptic vesicles arose from the spinal trigeminal nucleus and the lateral reticular nucleus. None were observed arising from the cuneate nucleus or dorsal root ganglion.

The mean diameters of vehicles from CuN, C2 DRG, LRN and Sp5 endings are 52.2 ± 10.5 nm (n = 484), 56.6 ± 8.6 nm (n = 642), 38.4 ± 6.9 (458) and 54.2 ± 9.9 nm (n = 530), respectively. The size of these vesicles from each area are significantly different (p < 0.0001, Post hoc Tukey test following One-way ANOVA), evaluated by the average diameters and distribution, except for those from cuneate and sP5 that the differences are slightly smaller (p < 0.005), but all others have different distribution of vesicles ranging from 40 to 80 nm (**Figure 12A**). Vesicles from CuN, C2 DRG and Sp5 are significantly larger compared to those from lateral reticular nucleus (p < 0.0001). They are, however, somewhat different in shape as characterized by long:short axis ratio (**Figure 12B**). All of the synaptic vesicles measured were slightly elongated.

**Figure 12.**
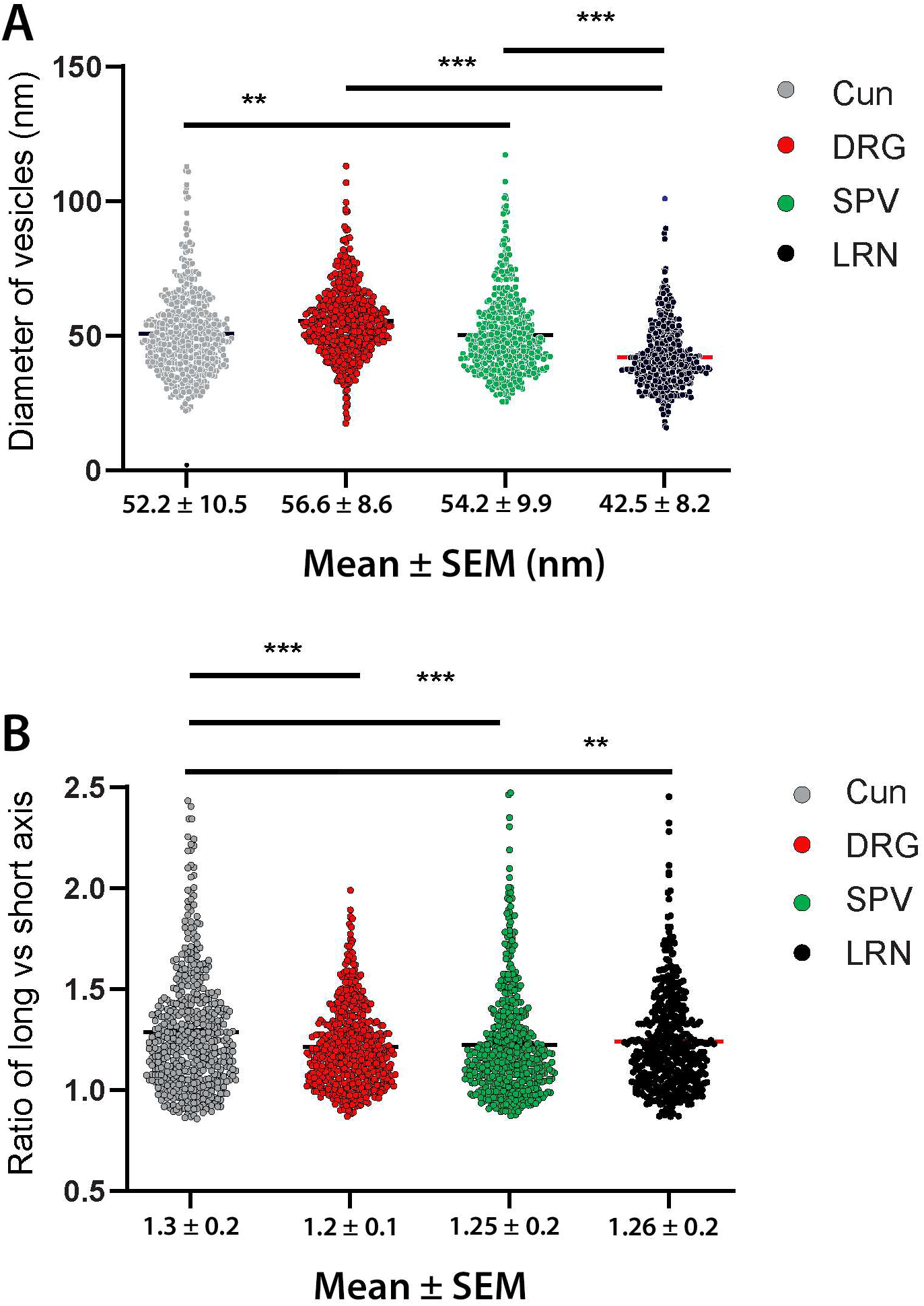
Distributions of synaptic vesicle diameters for terminals originating from the cuneate nucleus, C2 dorsal root ganglion (DRG), spinal trigeminal nucleus (sP5), and lateral reticular nucleus (LRN). Horizontal bars indicate pairwise comparisons between connected groups; significance levels are denoted as *** p < 0.0001 and ** p < 0.005 (Post hoc Tukey test following one-way ANOVA). **(A)** Synaptic vesicle diameters differ significantly among somatosensory structures. The difference between vesicles from the cuneate nucleus and sP5 is smaller but remains significant (p < 0.005). **(B)** Vesicle shape analysis reveals significant differences between vesicles from the cuneate nucleus and those from either C2 DRG or sP5 (p < 0.0001), whereas the difference between cuneate and LRN vesicles is modest but significant (p < 0.005). All other comparisons are not significant (p > 0.05).

## DISCUSSION

### The general synaptic architecture of mossy fibers in the GCD of cochlear nucleus

In general, a typical mossy fiber glomerulus contains mossy fiber, targeted dendrites or axons and associated microglia. We have examined some of the major medullary sensory inputs to the granule cell domain of the cochlear nucleus. These inputs appear primarily as mossy fibers converging onto the GCD of the dorsal cochlear nucleus. There are other times where the terminal is a bouton ending. The terminal distribution pattern for these inputs is remarkably similar. The CuN and Sp5 input endings are more prevalent and have more widespread distribution (Haenggeli et al., 2005; Wright & Ryugo, 1996), whereas the DRG and LRN have a moderate distribution (Zhan et al., 2006; Zhan & Ryugo, 2007). Emphasis on the size of the projection is premature at this time because the sizes of the dye injections were not uniform. Others have also shown non-auditory inputs to the GCD, including from the vestibular organ (Kevetter and Perachio, 1989; Gstoettner et al., 1992), nerve (Burian et al., 1989), and nuclei (Bukowska, 2002) but these publications do not comment about the size of the terminals nor specifically mention mossy fibers.

The mossy fibers typically have a large lobule structure, but boutons are much smaller. The boutons can arise independently from the projecting axon or from the mossy fibers of the same origin. The smaller size limits their postsynaptic targets. A mossy fiber can affect as many as 20 granule cells and each granule cell can receive potentially mossy fibers from the different modalities, such as C2 DRG, cuneate, spinal trigeminal and lateral reticular nucleus. Typically, for each mossy fiber or bouton, the targets are also innervated by a separate opposing excitatory ending. Since the small bouton derive from the same large mossy fibers, it may represent a transient state that mossy fibers undergo, a form of plasticity observed in climbing fibers in the cerebellum (Nishiyama, Fukaya, Watanabe, & Linden, 2007). Given that single inputs appear to have little power to elicit spikes, contributions from multiple and different inputs may be necessary to summate and activate the postsynaptic target.

A prominent feature of a mossy fiber is its large core of mitochondria. Mitochondria, the powerhouse of the cell are critical for the function of nerve terminals as the cycling of synaptic vesicle membrane requires a supply of ATP. The presynaptic mitochondria also take part in functions such as Ca^2+^ buffering and neurotransmitter synthesis. The core of mitochondria in a large mossy fiber occupies more than 20% of the total volume, which is identical across different modalities. Presynaptic mitochondria has been implicated to be closely related with the temporal pattern of neurotransmitter release (Brodin, Bakeeva, & Shupliakov, 1999). The mossy fibers may have a similar temporal drive to their postsynaptic targets. The aggregation and interconnection of mossy fiber mitochondria in a core, represents a feature as previously reported in other large synapses (Wimmer, Horstmann, Groh, & Kuner, 2006).

### The heterogeneity of presynaptic and postsynaptic structures of mossy fibers

Mossy fibers have diverse morphology, and they also differ in nature for their targets, thus forming distinct microcircuits. Somatosensory mossy fibers are incompletely surrounded by the dendrites of granule cells. As we have only investigated mossy fibers from one source at a time, inputs on the interested targets must be identified as an “unknown” source. In **Figure 2A**, the axon-axonal synapse implicates a presynaptic modulation on the C2 DRG input. In **Figure 2C**, the rare gap junction reminds us that there is still much to be learned about synaptic processing in the GCD. The granule cells and UBCs are the most common recipients of the mossy fiber inputs. Mossy fibers from the cuneate and spinal trigeminal nuclei were closely associated with the terminal dendritic claws of granule cells and formed many punctate synapses with these dendrites. Granule cells provide excitatory input to the inhibitory Golgi cells, which can in turn provide inhibitory feedback to more granule cells (Yaeger and Trussell, 2015). Golgi cells can spread inhibition to other granule cells and thereby influence the excitatory drive conveyed by granule cell parallel fibers to DCN pyramidal cells. Of note is that Golgi cells connect to each other via electrical synapses, such that a little input from a granule cell can greatly influence the level of inhibition in the GCD.

C2 DRG, dorsal column, LRN and sP5 all have inputs to the UBCs (**Figure 2, 5,7D, and 10**), whereas the spinal trigeminal nucleus form synapses with the Golgi cell body (**Figure 11**). Interestingly, the LRN can be either excitatory or inhibitory modulating postsynaptic targets (**Figures 7,8**). The most prominent mossy fiber synapses are excitatory and glutamate is the likely neurotransmitter (Petralia, Wang, Zhao, & Wenthold, 1996; Zeng et al., 2011), whereas glycine is an important inhibitory neurotransmitter in addition to GABA in the cochlear nucleus(Alibardi, 2003; Clements, Magnusson, & Beitz, 1990). Thus, each input contributes to a distinct microcircuitry. The specificity of the mossy fibers also displays unique vesicle properties for the sizes and densities, which may also have a unique chemical substrate for these synaptic vesicles, carrying unique information. These differences undoubtedly reflect functional differences yet to be determined.

All mossy fibers examined show multivesicular release site as there are several docked vesicles in one active zone. The vesicle-volume hypothesis posits that although the concentration of neurotransmitters within synaptic vesicles is maintained at a constant level, variations in the volume of these vesicles result in differences in the quantity of neurotransmitter released during an individual synaptic event. This discrepancy influences ’quantal size,’ which denotes the magnitude or amplitude of the response elicited in the postsynaptic neuron by the release of neurotransmitters from a single vesicle (Atwood & Karunanithi, 2002; Saviane & Silver, 2006). Thus there was a high variation of mEPSPs or mIPSPs when recording in the postsynaptic targets (Balakrishnan & Trussell, 2008). The large mossy fiber ending implicates a greater reserve pool of synaptic vesicles and easier vesicle priming and mobilization, or more accessible quantum availability. In the cerebellum, this great vesicle pool makes a prolongation of vesicle release possible when there is a longer-lasting Ca^2+^ entry caused by a longer duration of repetitive high frequency presynaptic activation. In the hippocampus, a high density of vesicles in mossy fibers, has been implicated that buffers calcium more effectively and thus prevents facilitation (Schikorski & Stevens, 1999). The large reserve vesicle pool of mossy fibers ensures high frequency transmission at the synapses (Atwood & Karunanithi, 2002; Saviane & Silver, 2006).

### C2 DRG neurons form substantial axo-axonic synapses in dorsal cochlear nucleus

The UBC appears to receive inputs exclusively from C2 DRG or cuneate nucleus (**Figure 2, 5 and 6**) (Zhan et al., 2006). Those mossy fibers originate from medium size DRG neurons whose neurotransmitters have been identified as glutamate, but possibly include substance P, somatostatin, 5-HT, dynorphin and enkephalin (Hao, Mata, Goins, Glorioso, & Fink, 2003; Kai-Kai, 1989; Price & Mudge, 1983). The inhibitory (axo-axonic) and excitatory (C2-GC or C2-UBC) projections in rats are consistent with the physiological responses in the principal neurons in cats (Kanold & Young, 2001; Koehler, Pradhan, Manis, & Shore, 2011). In contrast to the granule cells, the UBCs do not send axons to form parallel fibers targeting on fusiform cells, but they form synapses with other UBCs, cartwheel or granule cells (Dino & Mugnaini, 2008; Mugnaini, Sekerkova, & Martina, 2011; Oertel & Young, 2004). They have two distinct subtypes based on their physiological properties (Borges-Merjane & Trussell, 2015). The mossy fiber-UBC synaptic transmission is a slow one even though this is via fast AMPAR receptor activation, making it to be suitable for temporal coding (Lu, Balmer, Romero, & Trussell, 2017). The spinal trigeminal effects on principal neurons are diverse: excitation, excitation-inhibition or inhibition only, implicating a modulation of synaptic inhibition or excitation (Koehler et al., 2011; Shore, 2005). The effects from the non-trigeminal somatosensory inputs derived from C2 DRG on DCN neurons are mostly inhibitory, with some excitation and more inhibition at longer latencies (Davis, Miller, & Young, 1996; Kanold, Davis, & Young, 2011; Kanold & Manis, 2005; Kanold & Young, 2001; Young, Nelken, & Conley, 1995). The inhibition in the DCN circuitry has been suggested to participate neuronal responses of the principal neurons for the sensitivity to spectral notches (Bandyopadhyay, Reiss, & Young, 2007; Nelken & Young, 1994; Reiss & Young, 2005).

In addition to the direct inputs to the dorsal cochlear nucleus, a more widespread projections from dorsal root ganglion to the non-auditory brainstem have been reported, including major somatosensory nuclei: the cuneate nucleus, lateral reticular nucleus, vestibular and spinal trigeminal nuclei (Prihoda, Hiller, & Mayr, 1991; Zhang et al., 2015). All these latter nuclei project to the dorsal cochlear nucleus (Haenggeli et al., 2005; Vass et al., 1997; Wright & Ryugo, 1996; Zeng et al., 2011; Zhan & Ryugo, 2007). A cuneo-cochlear nucleus pathway originates mainly from the lateral part of the cuneate nucleus and the external cuneate nucleus that originates from C2 dorsal root ganglion, and this pathway has distinct effect on the principal neurons in DCN (Davis et al., 1996; Young et al., 1995). Cutaneous and proprioceptive trigeminal afferents of the head may supplement those of the lateral part of the cuneate nucleus which arise from neck and pinna muscle afferents as well as from cutaneous stretch receptors around the pinnae (Maslany, Crockett, & Egger, 1991; Miller, Richardson, Fibiger, & McLennan, 1975; Prihoda et al., 1991). Scalp projections from the C2 dorsal root ganglion produce a restricted terminal field in the medial sheet of the GCD along the posterior ventral cochlear nucleus and in the subpeduncular corner and lamina (Zhan et al., 2006). It is intriguingly puzzling whether these projections are the postsynaptic targets of C2 DRG that form axo-axonic synapses. Since the axo-axonic endings are indicative of synaptic inhibition (Hughes et al., 2012), presumably, they are an important mechanism of gating outputs from oncoming secondary somatosensory information by depressing action potentials arriving at the glomerulus, therefore, the contextually relevant information from different sources can be prioritized. This pre-synaptic machinery is particularly interesting, as it is plastic (Wefelmeyer, Cattaert, & Burrone, 2015). Despite somatosensory inputs to the granule cells are mostly excitatory, inhibition on granule cells from diverse sources is much more powerful that may sculpt the non-auditory inputs (Balakrishnan & Trussell, 2008).

More than two decade ago, the discovery of C2 DRG modulation of somatosensory inputs in the dorsal cochlear nucleus was proposed to be the underlying mechanism of modulation of clinical tinnitus (Levine, 1999). This hypothesis has been supported by more and more clinical studies on tinnitus or hyperacusis modulation (Delgado de la Serna et al., 2020; Demoen et al., 2023; Koning & Meulen, 2019; Michiels, van der Wal, et al., 2018; Spencer et al., 2022; van der Wal et al., 2022; Womack, Butts, & Dunning, 2022). Craniocervical stimulation and jaw manipulation following the perception of tinnitus provide circumstantial evidence implicating the involvement of somatosensory inputs. In addition, accumulating evidence suggests a role for unipolar brush cells in tinnitus (Bauer, Wisner, Baizer, & Brozoski, 2013; Brozoski, Brozoski, Wisner, & Bauer, 2017). These cells appear to play a critical role in polysensory integration, as they receive convergent input from the C2 dorsal root ganglion (DRG), cuneate nucleus, lateral reticular nucleus (LRN), and spinal trigeminal nucleus (sP5). This anatomical convergence places unipolar brush cells in a unique position to gate or modulate multisensory signals transmitted to principal neurons (Borges-Merjane & Trussell, 2015).. One other puzzling question is the axo-axonic mechanism from C2 DRG. Typically, the axo-axonic synapses in the central nervous system are GABAergic, that form a powerful inhibitory mechanism to attenuate the presynaptic activity (Fink et al., 2014; Zhang et al., 2015). Despite the axo-axonic synapses from C2 DRG appear to be symmetrical, they are from the branches of the putative excitatory C2 DRG axons. One hypothesis is that the axo-axonic synapses may be mediated by enkephalin or other neuromodulators as they are intensively distributed in the dorsal cochlear nucleus and dorsal root ganglions (Harlan, Shivers, Romano, Howells, & Pfaff, 1987; Zhang et al., 2015). Enkephalin and associated mu-receptors have appeared to be involved in pain modulation. This neurotransmitter is highly interesting as there is high coincidence that somatic tinnitus is associated a neck discomfort (Michiels, Ganz Sanchez, et al., 2018; Michiels, Van de Heyning, Truijen, Hallemans, & De Hertogh, 2016; Moller, 2007). Other neurotransmitters are also particularly interesting for their distribution in the dorsal cochlear nucleus and possible role for pre-synaptic transmission: 5-HT, somatostatin, and substance P (Tang & Trussell, 2015; Wynne & Robertson, 1997).

### Lateral reticular nucleus form synapses with resident cells in the dorsal cochlear nucleus

Despite its unique projection to dorsal cochlear nucleus, the function for LRN (**Figure 7 and 8**) in auditory processing is largely speculative (Zhan & Ryugo, 2007), specifically, it is not determined how LRN plays a role in hearing processing. One possible function is its contribution for proprioceptive inputs as the LRN receives projections from C3 and C4 spinal nerves (Kunzle, 1973). The peripheral processes of these cervical sensory nerves innervate neck, shoulder, and forelimb muscles, and their inputs signal movement of the head relative to the forelimbs and body (Ezure & Tanaka, 1997; Yates & Stocker, 1998). Stimulation of spinal nerve C2–C3 produced responses in the principal cells of the DCN, presumably conveying information about head-related transfer functions and sound directionality (Kanold & Young, 2001; Musicant, Chan, & Hind, 1990; Rice, May, Spirou, & Young, 1992). The LRN also sends mossy fibers to cerebellum and is considered a major pre-cerebellar center conveying a copy of information from the spinal system (Alstermark & Ekerot, 2013). We speculate that postural data (Flumerfelt, Hrycyshyn, & Kapogianis, 1982; Kunzle, 1973; Shokunbi, Hrycyshyn, & Flumerfelt, 1985, 1986) would complement information regarding pinna orientation (Kanold & Young, 2001; Rice et al., 1992), eye direction (Qvist, 1988; Qvist & Dietrichs, 1985, 1986), and body position with respect to gravity (Walberg, Dietrichs, & Nordby, 1985; Zhan & Ryugo, 2007). The integration of this collective sensory information could serve to stabilize and orient the body with respect to auditory, vestibular, and visual space. We predict that one target of the C2 DRG axo-axonic projection could be from LRN, so that complimentary somatosensory inputs can be integrated. Additionally, the LRN receives spinal input from visceral receptors that are thought to integrate cardiovascular and respiratory functions (Ezure & Tanaka, 1997; Macron, Marlot, & Duron, 1985). These studies have suggested autonomic influences on somatic motor activity (Yates & Stocker, 1998). Autonomic projections might also form the afferent limb of reflexes that suppresses auditory sensitivity to self-generated noises made by heartbeats, blood flow, and breathing (Singla, Dempsey, Warren, Enikolopov, & Sawtell, 2017). The suppression of internal noise is clearly different from the suppression of self-vocalizations or self-generated noise that involve the spinal trigeminal nucleus (Basura, Koehler, & Shore, 2012; Singla et al., 2017). Moreover, the LRN could have a role in vocalizations because of its involvement in respiratory activity (Jurgens, 2002). Vocalization is a complex behavior, the production of which involves the larynx, respiratory movements, and articulators. Vocal tract structure and respiration are influenced by the LRN; proper vocalization also requires feedback from the auditory system (Egnor, Iguina, & Hauser, 2006). Based on comparing vocal memories with vocalization feedback, the vocal motor system can compensate and make corrections to optimize communication. The dysfunction of this pathway may be the underlying mechanism for pulsatile tinnitus. Intensive investigation of LRN relevant circuitry in the DCN can make us understand its actual role in hearing processing or to clarify whether disturbance of this microcircuitry may cause phantom noise to be generated that is revealed as tinnitus. This line of research is particular relevant to pulsatile tinnitus that is not modulation by somatosensory manipulation.

### Functional significance of multimodal somatosensory integration in the dorsal cochlear nucleus

The DCN granule cell system, which receives information from the somatosensory and vestibular systems, would appear to have much of the necessary information about head movement and position to make this comparison. Descending projections that represent attention, affective state, or learned behaviors may serve to fine-tune this tracking system. As several somatosensory brainstem nuclei are involved in initial processing, spatial and motion information are carried by these inputs to dorsal cochlear nucleus. Similar strategy has been detected in vision perception (Meyer, Poort, O’Keefe, Sahani, & Linden, 2018). The visual system should also play some role in this process, so it would be practical to search for visual inputs to the auditory pathway, and lateral reticular nucleus may serve as this role (Qvist, 1988; Qvist & Dietrichs, 1985). The somatosensory integration along with binaural mechanisms by using timing differences would work together to fulfill for encoding of sound location (Carr, 1993; Oertel, 1999).

This produces persistent, use-dependent alterations in synaptic strength postsynaptically as a decrease in the synaptic current evoked by a fixed delivery of transmitter. Mossy fibers have numerous active zones which may also contribute an increase in the number of functional synaptic contacts for other types of synaptic plasticity and are able to integrate granule cell spiking activity over a broad range of frequencies (Salin, Scanziani, Malenka, & Nicoll, 1996). Our findings support the idea that synapses are not alike, although the mossy fibers from different sources are similar based on the profiles. They are likely different functionally, thus carry unique signals pertaining to the specific mode. Moreover, given the diverse sources of input to the granule cell system, these neurons may integrate signals from different brain regions and sensory modalities, providing a sensory prediction (Sawtell, 2010). The UBCs were implicated to amplify signals, as they send axons to granule cells and other UBCs, this essentially make the feed-forward synaptic transmission stronger (Mugnaini et al., 2011). The Golgi cells have been implicated to provide a feedback gain control by inhibiting granule cells (Cesana et al., 2013).

However, it argues that the MF-GoC-GrC circuit in cerebellum is a feed-forward mechanism used to conserve temporal precision (Kanichay & Silver, 2008). The spiking of this cell is modulated by M-receptors in the cochlear nucleus, suggesting cholinergic involvement in possible arousal responses (Irie, Fukui, & Ohmori, 2006; Mellott, Motts, & Schofield, 2011). The diverse multimodal inputs to the dorsal cochlear nucleus and comprehensive interactive connections among the C2 DRG, cuneate, lateral reticular, spinal trigeminal inputs and local resident cells make it a suitable candidate to extract temporal information(Huson, Newman, & Regehr, 2023). As this structure is intrinsically associated with auditory system, a dysfunctional or maladaptive modification of the somatosensory inputs may lead to generate internal “noise” or somatic tinnitus (Eggermont, 2007; Kaltenbach, Zhang, & Finlayson, 2005; Wu, Stefanescu, Martel, & Shore, 2016).

## CONCLUSIONS

Sensory information regarding head, neck and pinna position are conveyed by skin stretch receptors and muscle proprioceptors to provide information relevant to sound source separation for the central auditory system. The information converges directly or indirectly onto granule cell dendrites in the cochlear nucleus, forming a unique system of microcircuits. In the granule cell domain, afferent mossy fibers also contact excitatory or inhibitory UBCs that send feed-forward input to additional granule cells or other UBCs. The axonal inputs may be modulated by inputs from other modalities. The fibers from the spinal trigeminal nucleus and lateral reticular nucleus are special because of their direct inputs, excitatory or inhibitory, on the somata of inhibitory Golgi cells. The LRN fibers also provide distinct inhibitory inputs, presumably to attenuate inputs from other modalities. The synaptic contribution from these various circuits determines the multimodal signals delivered to the projecting cells of the dorsal cochlear nucleus that integrates somatosensory, vestibular and visual information with auditory cues for localizing and separating the positions of sounds in space.

## ACKNOWLEDGEMENTS

This work was supported by National Institutes of Health (RO1 DC04395), the National Health and Medical Research Council of Australia (1080652 and 1081478), and donations from Alan and Lynne Rydge, Haydn and Sue Daw, Christina Kennedy, and Charlene and Graham Bradley to DKR; and in part by the *Les Paul* Foundation (via Hearing Heath Foundation), Karen Toffler Charitable Trust and the HUCOM bridge fund to XZ. We thank Tan Pongstaphone and Ping Wang, for invaluable technical assistance.

## Abbreviation

C2 DRG,: second cervical dorsal root ganglion
CuN,: cuneate nucleus
DCN,: dorsal cochlear nucleus
EPSP,: excitatory postsynaptic potential
GCD,: granule cell domain
PSD,: postsynaptic density
Sp5,: spinal trigeminal nucleus
UBC,: unipolar brush cel

## References

1. Alibardi, L. (2003). Ultrastructural immunocytochemistry for glycine in neurons of the dorsal cochlear nucleus of the guinea pig. J Submicrosc Cytol Pathol, 35(4), 373–387.

2. Alstermark, B., & Ekerot, C. F. (2013). The lateral reticular nucleus: a precerebellar centre providing the cerebellum with overview and integration of motor functions at systems level. A new hypothesis. J Physiol, 591(22), 5453–5458. doi:10.1113/jphysiol.2013.256669

3. Atwood, H. L., & Karunanithi, S. (2002). Diversification of synaptic strength: presynaptic elements. Nat Rev Neurosci, 3(7), 497–516.

4. Balakrishnan, V., & Trussell, L. O. (2008). Synaptic inputs to granule cells of the dorsal cochlear nucleus. J Neurophysiol, 99(1), 208–219.

5. Bandyopadhyay, S., Reiss, L. A., & Young, E. D. (2007). Receptive field for dorsal cochlear nucleus neurons at multiple sound levels. J Neurophysiol, 98(6), 3505–3515.

6. Basura, G. J., Koehler, S. D., & Shore, S. E. (2012). Multi-sensory integration in brainstem and auditory cortex. Brain Res, 1485, 95–107. doi:10.1016/j.brainres.2012.08.037

7. Bauer, C. A., Wisner, K. W., Baizer, J. S., & Brozoski, T. J. (2013). Tinnitus, unipolar brush cells, and cerebellar glutamatergic function in an animal model. PLoS One, 8(6), e64726. doi:10.1371/journal.pone.0064726

8. Borges-Merjane, C., & Trussell, L. O. (2015). ON and OFF unipolar brush cells transform multisensory inputs to the auditory system. Neuron, 85(5), 1029–1042. doi:10.1016/j.neuron.2015.02.009

9. Brodin, L., Bakeeva, L., & Shupliakov, O. (1999). Presynaptic mitochondria and the temporal pattern of neurotransmitter release. Philos Trans R Soc Lond B Biol Sci, 354(1381), 365–372.

10. Brozoski, T., Brozoski, D., Wisner, K., & Bauer, C. (2017). Chronic tinnitus and unipolar brush cell alterations in the cerebellum and dorsal cochlear nucleus. Hear Res, 350, 139–151. doi:10.1016/j.heares.2017.04.016

11. Carr, C. E. (1993). Processing of temporal information in the brain. Annu Rev Neurosci, 16, 223–243. doi:10.1146/annurev.ne.16.030193.001255

12. Cesana, E., Pietrajtis, K., Bidoret, C., Isope, P., D’Angelo, E., Dieudonne, S., & Forti, L. (2013). Granule cell ascending axon excitatory synapses onto Golgi cells implement a potent feedback circuit in the cerebellar granular layer. J Neurosci, 33(30), 12430–12446. doi:10.1523/JNEUROSCI.4897-11.201333/30/12430 [pii]

13. Clements, J. R., Magnusson, K. R., & Beitz, A. J. (1990). Ultrastructural description of glutamate-, aspartate-, taurine-, and glycine-like immunoreactive terminals from five rat brain regions. J Electron Microsc Tech, 15(1), 49–66. doi:10.1002/jemt.1060150106

14. Davis, K. A., Miller, R. L., & Young, E. D. (1996). Effects of somatosensory and parallel-fiber stimulation on neurons in dorsal cochlear nucleus. J Neurophysiol, 76(5), 3012–3024.

15. Delgado de la Serna, P., Plaza-Manzano, G., Cleland, J., Fernandez-de-Las-Penas, C., Martin-Casas, P., & Diaz-Arribas, M. J. (2020). Effects of Cervico-Mandibular Manual Therapy in Patients with Temporomandibular Pain Disorders and Associated Somatic Tinnitus: A Randomized Clinical Trial. Pain Med, 21(3), 613–624. doi:10.1093/pm/pnz278

16. Demoen, S., Michiels, S., Gilles, A., Vermeersch, H., Joossen, I., Vanderveken, O. M., . . . Jacquemin, L. (2023). Pilot study on the role of somatic modulation in hyperacusis. Eur Arch Otorhinolaryngol, 280(3), 1425–1435. doi:10.1007/s00405-022-07695-y

17. Dino, M. R., & Mugnaini, E. (2000). Postsynaptic actin filaments at the giant mossy fiber-unipolar brush cell synapse. Synapse, 38(4), 499–510.

18. Dino, M. R., & Mugnaini, E. (2008). Distribution and phenotypes of unipolar brush cells in relation to the granule cell system of the rat cochlear nucleus. Neuroscience, 154(1), 29–50. doi:S0306-4522(08)00108-5 [pii] 10.1016/j.neuroscience.2008.01.035

19. Eggermont, J. J. (2007). Pathophysiology of tinnitus. Prog Brain Res, 166, 19–35. doi:S0079-6123(07)66002-6 [pii] 10.1016/S0079-6123(07)66002-6

20. Egnor, S. E., Iguina, C. G., & Hauser, M. D. (2006). Perturbation of auditory feedback causes systematic perturbation in vocal structure in adult cotton-top tamarins. J Exp Biol, 209(Pt 18), 3652–3663. doi:10.1242/jeb.02420

21. Ezure, K., & Tanaka, I. (1997). Convergence of central respiratory and locomotor rhythms onto single neurons of the lateral reticular nucleus. Exp Brain Res, 113(2), 230–242. doi:10.1007/BF02450321

22. Fink, A. J., Croce, K. R., Huang, Z. J., Abbott, L. F., Jessell, T. M., & Azim, E. (2014). Presynaptic inhibition of spinal sensory feedback ensures smooth movement. Nature, 509(7498), 43–48. doi:10.1038/nature13276

23. Floris, A., Dino, M., Jacobowitz, D. M., & Mugnaini, E. (1994). The unipolar brush cells of the rat cerebellar cortex and cochlear nucleus are calretinin-positive: a study by light and electron microscopic immunocytochemistry. Anat Embryol (Berl), 189(6), 495–520.

24. Flumerfelt, B. A., Hrycyshyn, A. W., & Kapogianis, E. M. (1982). Spinal projections to the lateral reticular nucleus in the rat. Anat Embryol (Berl), 165(3), 345–359.

25. Haenggeli, C. A., Pongstaporn, T., Doucet, J. R., & Ryugo, D. K. (2005). Projections from the spinal trigeminal nucleus to the cochlear nucleus in the rat. J Comp Neurol, 484(2), 191–205.

26. Hao, S., Mata, M., Goins, W., Glorioso, J. C., & Fink, D. J. (2003). Transgene-mediated enkephalin release enhances the effect of morphine and evades tolerance to produce a sustained antiallodynic effect in neuropathic pain. Pain, 102(1-2), 135–142.

27. Harlan, R. E., Shivers, B. D., Romano, G. J., Howells, R. D., & Pfaff, D. W. (1987). Localization of preproenkephalin mRNA in the rat brain and spinal cord by in situ hybridization. J Comp Neurol, 258(2), 159–184. doi:10.1002/cne.902580202

28. Hughes, D. I., Sikander, S., Kinnon, C. M., Boyle, K. A., Watanabe, M., Callister, R. J., & Graham, B. A. (2012). Morphological, neurochemical and electrophysiological features of parvalbumin-expressing cells: a likely source of axo-axonic inputs in the mouse spinal dorsal horn. J Physiol, 590(16), 3927–3951. doi:10.1113/jphysiol.2012.235655

29. Huson, V., Newman, L., & Regehr, W. G. (2023). A Continuum of Response Properties across the Population of Unipolar Brush Cells in the Dorsal Cochlear Nucleus. J Neurosci, 43(34), 6035–6045. doi:10.1523/JNEUROSCI.0873-23.2023

30. Irie, T., Fukui, I., & Ohmori, H. (2006). Activation of GIRK channels by muscarinic receptors and group II metabotropic glutamate receptors suppresses Golgi cell activity in the cochlear nucleus of mice. J Neurophysiol, 96(5), 2633–2644. doi:00396.2006 [pii] 10.1152/jn.00396.2006

31. Jurgens, U. (2002). Neural pathways underlying vocal control. Neurosci Biobehav Rev, 26(2), 235–258.

32. Kai-Kai, M. A. (1989). Cytochemistry of the trigeminal and dorsal root ganglia and spinal cord of the rat. Comp Biochem Physiol A Comp Physiol, 93(1), 183–193.

33. Kaltenbach, J. A., Zhang, J., & Finlayson, P. (2005). Tinnitus as a plastic phenomenon and its possible neural underpinnings in the dorsal cochlear nucleus. Hear Res, 206(1-2), 200–226.

34. Kanichay, R. T., & Silver, R. A. (2008). Synaptic and cellular properties of the feedforward inhibitory circuit within the input layer of the cerebellar cortex. J Neurosci, 28(36), 8955–8967. doi:10.1523/JNEUROSCI.5469-07.200828/36/8955 [pii]

35. Kanold, P. O., Davis, K. A., & Young, E. D. (2011). Somatosensory context alters auditory responses in the cochlear nucleus. J Neurophysiol, 105(3), 1063–1070. doi:10.1152/jn.00807.2010jn.00807.2010 [pii]

36. Kanold, P. O., & Manis, P. B. (2005). Encoding the timing of inhibitory inputs. J Neurophysiol, 93(5), 2887–2897.

37. Kanold, P. O., & Young, E. D. (2001). Proprioceptive information from the pinna provides somatosensory input to cat dorsal cochlear nucleus. J Neurosci, 21(19), 7848–7858.

38. Koehler, S. D., Pradhan, S., Manis, P. B., & Shore, S. E. (2011). Somatosensory inputs modify auditory spike timing in dorsal cochlear nucleus principal cells. Eur J Neurosci, 33(3), 409–420. doi:10.1111/j.1460-9568.2010.07547.x

39. Koning, H. M., & Meulen, B. C. T. (2019). Pulsed radiofrequency of C2 dorsal root ganglion in patients with tinnitus. Int Tinnitus J, 23(2), 91–96. doi:10.5935/0946-5448.20190016

40. Kunzle, H. (1973). The topographic organization of spinal afferents to the lateral reticular nucleus of the cat. J Comp Neurol, 149(1), 103–115.

41. Levine, R. A. (1999). Somatic (craniocervical) tinnitus and the dorsal cochlear nucleus hypothesis. Am J Otolaryngol, 20(6), 351–362.

42. Lu, H. W., Balmer, T. S., Romero, G. E., & Trussell, L. O. (2017). Slow AMPAR Synaptic Transmission Is Determined by Stargazin and Glutamate Transporters. Neuron, 96(1), 73–80 e74. doi:10.1016/j.neuron.2017.08.043

43. Macron, J. M., Marlot, D., & Duron, B. (1985). Phrenic afferent input to the lateral medullary reticular formation of the cat. Respir Physiol, 59(2), 155–167. doi:10.1016/0034-5687(85)90004-0

44. Maslany, S., Crockett, D. P., & Egger, M. D. (1991). Somatotopic organization of the dorsal column nuclei in the rat: transganglionic labelling with B-HRP and WGA-HRP. Brain Res, 564(1), 56–65.

45. Mellott, J. G., Motts, S. D., & Schofield, B. R. (2011). Multiple origins of cholinergic innervation of the cochlear nucleus. Neuroscience, 180, 138–147. doi:10.1016/j.neuroscience.2011.02.010S0306-4522(11)00151-5 [pii]

46. Meyer, A. F., Poort, J., O’Keefe, J., Sahani, M., & Linden, J. F. (2018). A Head-Mounted Camera System Integrates Detailed Behavioral Monitoring with Multichannel Electrophysiology in Freely Moving Mice. Neuron, 100(1), 46–60 e47. doi:10.1016/j.neuron.2018.09.020

47. Michiels, S., Ganz Sanchez, T., Oron, Y., Gilles, A., Haider, H. F., Erlandsson, S., … Hall, D. A. (2018). Diagnostic Criteria for Somatosensory Tinnitus: A Delphi Process and Face-to-Face Meeting to Establish Consensus. Trends Hear, 22, 2331216518796403. doi:10.1177/2331216518796403

48. Michiels, S., Van de Heyning, P., Truijen, S., Hallemans, A., & De Hertogh, W. (2016). Does multi-modal cervical physical therapy improve tinnitus in patients with cervicogenic somatic tinnitus? Man Ther, 26, 125–131. doi:10.1016/j.math.2016.08.005

49. Michiels, S., van der Wal, A. C., Nieste, E., Van de Heyning, P., Braem, M., Visscher, C., … De Hertogh, W. (2018). Conservative therapy for the treatment of patients with somatic tinnitus attributed to temporomandibular dysfunction: study protocol of a randomised controlled trial. Trials, 19(1), 554. doi:10.1186/s13063-018-2903-1

50. Miller, J. J., Richardson, T. L., Fibiger, H. C., & McLennan, H. (1975). Anatomical and electrophysiological identification of a projection from the mesencephalic raphe to the caudate-putamen in the rat. Brain Res, 97(1), 133–136.

51. Moller, A. R. (2007). Tinnitus and pain. Prog Brain Res, 166, 47–53. doi:S0079-6123(07)66004-X [pii] 10.1016/S0079-6123(07)66004-X

52. Mugnaini, E., Dino, M. R., & Jaarsma, D. (1997). The unipolar brush cells of the mammalian cerebellum and cochlear nucleus: cytology and microcircuitry. Prog Brain Res, 114, 131–150.

53. Mugnaini, E., & Floris, A. (1994). The unipolar brush cell: a neglected neuron of the mammalian cerebellar cortex. J Comp Neurol, 339(2), 174–180.

54. Mugnaini, E., Sekerkova, G., & Martina, M. (2011). The unipolar brush cell: a remarkable neuron finally receiving deserved attention. Brain Res Rev, 66(1-2), 220–245. doi:10.1016/j.brainresrev.2010.10.001 S0165-0173(10)00112-8 [pii]

55. Musicant, A. D., Chan, J. C., & Hind, J. E. (1990). Direction-dependent spectral properties of cat external ear: new data and cross-species comparisons. J Acoust Soc Am, 87(2), 757–781.

56. Nelken, I., & Young, E. D. (1994). Two separate inhibitory mechanisms shape the responses of dorsal cochlear nucleus type IV units to narrowband and wideband stimuli. J Neurophysiol, 71(6), 2446–2462.

57. Nishiyama, H., Fukaya, M., Watanabe, M., & Linden, D. J. (2007). Axonal motility and its modulation by activity are branch-type specific in the intact adult cerebellum. Neuron, 56(3), 472–487.

58. Oertel, D. (1999). The role of timing in the brain stem auditory nuclei of vertebrates. Annu Rev Physiol, 61, 497–519. doi:10.1146/annurev.physiol.61.1.497

59. Oertel, D., & Young, E. D. (2004). What’s a cerebellar circuit doing in the auditory system? Trends Neurosci, 27(2), 104–110.

60. Petralia, R. S., Wang, Y. X., Zhao, H. M., & Wenthold, R. J. (1996). Ionotropic and metabotropic glutamate receptors show unique postsynaptic, presynaptic, and glial localizations in the dorsal cochlear nucleus. J Comp Neurol, 372(3), 356–383. doi:10.1002/(SICI)1096-9861(19960826)372:3<356::AID-CNE3>3.0.CO;2-1

61. Price, J., & Mudge, A. W. (1983). A subpopulation of rat dorsal root ganglion neurones is catecholaminergic. Nature, 301(5897), 241–243.

62. Prihoda, M., Hiller, M. S., & Mayr, R. (1991). Central projections of cervical primary afferent fibers in the guinea pig: an HRP and WGA/HRP tracer study. J Comp Neurol, 308(3), 418–431. doi:10.1002/cne.903080309

63. Qvist, H. (1988). Afferents to the lateral reticular nucleus from the oculomotor region. II. The oculomotor nucleus, the interstitial nucleus of Cajal and the nucleus of the posterior commissure. Anat Embryol (Berl), 177(3), 277–283.

64. Qvist, H., & Dietrichs, E. (1985). The projection from the superior colliculus to the lateral reticular nucleus in the cat as studied with retrograde transport of WGA-HRP. Anat Embryol (Berl), 173(2), 269–274.

65. Qvist, H., & Dietrichs, E. (1986). Afferents to the lateral reticular nucleus from the oculomotor region. I. The Edinger-Westphal nucleus. Anat Embryol (Berl), 175(2), 261–269.

66. Reiss, L. A., & Young, E. D. (2005). Spectral edge sensitivity in neural circuits of the dorsal cochlear nucleus. J Neurosci, 25(14), 3680–3691.

67. Rice, J. J., May, B. J., Spirou, G. A., & Young, E. D. (1992). Pinna-based spectral cues for sound localization in cat. Hear Res, 58(2), 132–152.

68. Rizzoli, S. O., & Betz, W. J. (2005). Synaptic vesicle pools. Nat Rev Neurosci, 6(1), 57–69.

69. Salin, P. A., Scanziani, M., Malenka, R. C., & Nicoll, R. A. (1996). Distinct short-term plasticity at two excitatory synapses in the hippocampus. Proc Natl Acad Sci U S A, 93(23), 13304–13309.

70. Saviane, C., & Silver, R. A. (2006). Fast vesicle reloading and a large pool sustain high bandwidth transmission at a central synapse. Nature, 439(7079), 983–987.

71. Sawtell, N. B. (2010). Multimodal integration in granule cells as a basis for associative plasticity and sensory prediction in a cerebellum-like circuit. Neuron, 66(4), 573–584. doi:S0896-6273(10)00289-8 [pii] 10.1016/j.neuron.2010.04.018

72. Schikorski, T., & Stevens, C. F. (1999). Quantitative fine-structural analysis of olfactory cortical synapses. Proc Natl Acad Sci U S A, 96(7), 4107–4112.

73. Shokunbi, M. T., Hrycyshyn, A. W., & Flumerfelt, B. A. (1985). Spinal projections to the lateral reticular nucleus in the rat: a retrograde labelling study using horseradish peroxidase. J Comp Neurol, 239(2), 216–226.

74. Shokunbi, M. T., Hrycyshyn, A. W., & Flumerfelt, B. A. (1986). A horseradish peroxidase study of the rubral and cortical afferents to the lateral reticular nucleus in the rat. J Comp Neurol, 248(3), 441–454. doi:10.1002/cne.902480312

75. Shore, S. E. (2005). Multisensory integration in the dorsal cochlear nucleus: unit responses to acoustic and trigeminal ganglion stimulation. Eur J Neurosci, 21(12), 3334–3348.

76. Singla, S., Dempsey, C., Warren, R., Enikolopov, A. G., & Sawtell, N. B. (2017). A cerebellum-like circuit in the auditory system cancels responses to self-generated sounds. Nat Neurosci, 20(7), 943–950. doi:10.1038/nn.4567

77. Spencer, S., Mielczarek, M., Olszewski, J., Sereda, M., Joossen, I., Vermeersch, H., … Michiels, S. (2022). Effectiveness of bimodal auditory and electrical stimulation in patients with tinnitus: A feasibility study. Front Neurosci, 16, 971633. doi:10.3389/fnins.2022.971633

78. Tang, Z. Q., & Trussell, L. O. (2015). Serotonergic regulation of excitability of principal cells of the dorsal cochlear nucleus. J Neurosci, 35(11), 4540–4551. doi:10.1523/JNEUROSCI.4825-14.2015

79. van der Wal, A., Michiels, S., Van de Heyning, P., Gilles, A., Jacquemin, L., Van Rompaey, V., … De Hertogh, W. (2022). Reduction of Somatic Tinnitus Severity is Mediated by Improvement of Temporomandibular Disorders. Otol Neurotol, 43(3), e309–e315. doi:10.1097/MAO.0000000000003446

80. Vass, Z., Shore, S. E., Nuttall, A. L., Jancso, G., Brechtelsbauer, P. B., & Miller, J. M. (1997). Trigeminal ganglion innervation of the cochlea--a retrograde transport study. Neuroscience, 79(2), 605–615. doi:S0306452296006410 [pii]

81. Walberg, F., Dietrichs, E., & Nordby, T. (1985). On the projections from the vestibular and perihypoglossal nuclei to the spinal trigeminal and lateral reticular nuclei in the cat. Brain Res, 333(1), 123–130.

82. Wefelmeyer, W., Cattaert, D., & Burrone, J. (2015). Activity-dependent mismatch between axo-axonic synapses and the axon initial segment controls neuronal output. Proc Natl Acad Sci U S A, 112(31), 9757–9762. doi:10.1073/pnas.1502902112

83. Wimmer, V. C., Horstmann, H., Groh, A., & Kuner, T. (2006). Donut-like topology of synaptic vesicles with a central cluster of mitochondria wrapped into membrane protrusions: a novel structure-function module of the adult calyx of Held. J Neurosci, 26(1), 109–116.

84. Womack, A., Butts, R., & Dunning, J. (2022). Dry needling as a novel intervention for cervicogenic somatosensory tinnitus: a case study. Physiother Theory Pract, 38(9), 1319–1327. doi:10.1080/09593985.2020.1825579

85. Wouterlood, F. G., & Mugnaini, E. (1984). Cartwheel neurons of the dorsal cochlear nucleus: a Golgi-electron microscopic study in rat. J Comp Neurol, 227(1), 136–157. doi:10.1002/cne.902270114

86. Wouterlood, F. G., Mugnaini, E., Osen, K. K., & Dahl, A. L. (1984). Stellate neurons in rat dorsal cochlear nucleus studies with combined Golgi impregnation and electron microscopy: synaptic connections and mutual coupling by gap junctions. J Neurocytol, 13(4), 639–664.

87. Wright, D. D., & Ryugo, D. K. (1996). Mossy fiber projections from the cuneate nucleus to the cochlear nucleus in the rat. J Comp Neurol, 365(1), 159–172.

88. Wu, C., Stefanescu, R. A., Martel, D. T., & Shore, S. E. (2016). Tinnitus: Maladaptive auditory-somatosensory plasticity. Hear Res, 334, 20–29. doi:10.1016/j.heares.2015.06.005

89. Wynne, B., & Robertson, D. (1997). Somatostatin and substance P-like immunoreactivity in the auditory brainstem of the adult rat. J Chem Neuroanat, 12(4), 259–266.

90. Yaeger, D. B., & Trussell, L. O. (2015). Single granule cells excite Golgi cells and evoke feedback inhibition in the cochlear nucleus. J Neurosci, 35(11), 4741–4750. doi:10.1523/JNEUROSCI.3665-14.2015

91. Yates, B. J., & Stocker, S. D. (1998). Integration of somatic and visceral inputs by the brainstem: functional considerations. Exp Brain Res, 119(3), 269–275.

92. Young, E. D., Nelken, I., & Conley, R. A. (1995). Somatosensory effects on neurons in dorsal cochlear nucleus. J Neurophysiol, 73(2), 743–765.

93. Zeng, C., Shroff, H., & Shore, S. E. (2011). Cuneate and spinal trigeminal nucleus projections to the cochlear nucleus are differentially associated with vesicular glutamate transporter-2. Neuroscience, 176, 142–151. doi:S0306-4522(10)01587-3 [pii] 10.1016/j.neuroscience.2010.12.010

94. Zhan, X., Pongstaporn, T., & Ryugo, D. K. (2006). Projections of the second cervical dorsal root ganglion to the cochlear nucleus in rats. J Comp Neurol, 496(3), 335–348.

95. Zhan, X., & Ryugo, D. K. (2007). Projections of the lateral reticular nucleus to the cochlear nucleus in rats. J Comp Neurol, 504(5), 583–598.

96. Zhang, Y., Zhao, S., Rodriguez, E., Takatoh, J., Han, B. X., Zhou, X., & Wang, F. (2015). Identifying local and descending inputs for primary sensory neurons. J Clin Invest, 125(10), 3782–3794. doi:10.1172/JCI81156

